# Persistent Microglial HIV Infection Drives Neuroinflammation Despite Viral Suppression: Insights from Rapid Postmortem Brain Biopsies

**DOI:** 10.64898/2026.05.20.726582

**Authors:** M. Nühn Marieke, Nadia Sabet, Nanouk Zuidmeer, C.J. van Abeelen Kirsten, E. Hermans Lucas, J. Schipper Pauline, Raphael Kübler, E. Basson Adriaan, Tanvier Omar, Ebrahim Variava, A. Martinson Neil, Stefanie Giacopazzi, D. F. Venter Willem, J. Muraro Mauro, M. J. Wensing Annemarie, D. de Witte Lot, A. Papathanasopoulos Maria, Monique Nijhuis, Jori Symons

## Abstract

Despite suppressive antiretroviral therapy (ART), HIV persists in the central nervous system (CNS) and contributes to HIV-associated neurocognitive disorder (HAND), but cell type–specific effects remain poorly defined. Using fluorescence-activated nuclei sorting of postmortem brain tissue of aviremic and viremic deceased people with HIV (DPWH) and HIV-negative individuals, we quantified the size of the HIV CNS reservoir and transcriptional alterations. Microglia were identified as the dominant CNS reservoir, harboring 10³–10⁴ HIV DNA copies per million cells by ddPCR-LTR assay in both aviremic and viremic DPWH. Bulk RNA-sequencing revealed immune pathway upregulation specifically in microglia, and downregulation of synaptic and homeostatic pathways across cell-types in viremic compared to aviremic individuals. ART partially mitigated microglial transcriptional dysregulation, but transcriptional profiles did not restore profiles to HIV-negative levels. Notably, persistent microglial infection was associated with transcriptional changes in other cell-types, underscoring microglia as a key therapeutical target for CNS-directed HIV cure strategies.

## Introduction

Despite viral suppression by antiretroviral therapy (ART) human immunodeficiency virus (HIV) can persist in various tissues[1, 2], with the central nervous system (CNS) serving as an anatomical reservoir[3, 4]. HIV persistence can contribute to neurological complications, termed HIV associated neurocognitive disorders (HAND). These disorders cause cognitive and behavioral impairments that can significantly impact the quality of life. Although ART has reduced the frequency and severity of HAND, it can still be observed in people on suppressive ART, underscoring the chronic impact of HIV on the CNS[5, 6]. Systemic and neuroinflammation, along with the direct and indirect toxic effects of viral products, are believed to contribute to neuronal dysfunction or damage[7–9]. However, the precise effects of HIV on different brain cells and the influence of ART on CNS inflammation and functioning remain poorly understood.

The detection of HIV transcripts and proviral DNA in brains of deceased people with HIV (DPWH) on ART[10–13], strongly indicates that HIV persists in the CNS and contributes to the development of neuroinflammation. In general, HIV infection results in the production of viral transcripts and viral proteins, evoking a pro-inflammatory cytokine response, fueling chronic systemic inflammation[14]. These viral proteins and pro-inflammatory cytokines are known to be neurotoxic[15, 16], which is reflected by elevated levels of neopterin in cerebrospinal fluid (CSF)[17] and neurofilament light chain protein (NfL) in both CSF and blood[18]. Both serve as biomarkers of neuroinflammation and neurodegeneration, respectively, in some people with HIV (PWH). These findings suggest that HIV persistence drives neuroinflammation and neurodegeneration. Although ART effectively suppresses viral replication and systemic inflammation[19], low-level systemic viral transcription, protein production and inflammation can persist despite treatment[20–22]. Despite good CNS penetration of many ART regimens, HIV transcripts have also been detected in the brains of both aviremic (undetectable viral load) and viremic (detectable viral load) individuals[13], likely due to viral production rather than replication[23]. Ultimately, the impact of viral supression and HIV persistence on CNS neuroinflammation remains unclear.

Proviral HIV DNA of both defective and replication-competent viruses are predominantly found in microglia[11, 13, 24–26], which are widely recognized as the primary HIV reservoir in the CNS. Nevertheless, other cell types, including astrocytes[24, 27] and infiltrating T cells[28, 29] have also been proposed as potential reservoirs. However, the size and distribution of the HIV proviral reservoir across these cell types remain poorly defined. HIV infection of microglia results in a pro-inflammatory state and contributes to neuroinflammation and neuronal cell death[30–32]. Moreover, other resident brain cells, including neurons, oligodendrocytes, and astrocytes, which are essential for maintaining normal brain function, can also be indirectly damaged by viral products or the inflammatory environment even in the absence of direct infection [32–34]. Recent studies found evidence of increased immune activity and reduced homeostatic and metabolic functions in both microglia and astrocytes of DPWH compared to HIV-negative individuals[35, 36]. However, the specific impact of viral supression on these changes remains unclear. Therefore, it is critical that the cell type-specific effects of HIV on brain function in aviremic and viremic PWH are examined.

Studying CNS reservoirs is challenging due to the limited availability of high-quality brain tissues with detailed clinical and histopathological data to correlate. In contrast, this study uniquely leveraged rapidly obtained postmortem brain tissue of both aviremic (n=7) and viremic individuals (n=4), and HIV-negative individuals (n=3) with extensively documented antemortem data, postmortem viral loads measurements, and clinical and histological pathology including cause of death reviews, enabling a more nuanced understanding of HIV reservoir dynamics. Additionally, CNS reservoir studies face challenges in isolating specific cell populations due to the heterogeneity of cells which are heavily interconnected in brain tissue[37]. However, fluorescence-activated nuclei sorting (FANS) allows for isolation and analysis of cell-type specific nuclear fractions[37, 38]. Using FANS, this study uniquely investigates the cell-type specific impact of viral suppression in brains of DPWH. We aimed to uncover cell-type specific effects of HIV persistence, its contribution to neuroinflammation, and its impact on brain functioning in both aviremic and viremic individuals. Gaining a deeper understanding of these mechanisms is crucial for improving treatment of HIV infection and developing HIV cure strategies that are both effective and safe in the CNS.

## Results

### Participant characteristics

Postmortem brain biopsies were taken from 14 individuals, including 11 DPWH (H1-11) and 3 HIV negative individuals (N1-3). H1, H3, H8 and H11 were viremic, with plasma median viral loads of 55,579 HIV-RNA copies/mL, ranging from 1,840-158,000 HIV-RNA copies/mL, while the other PDWH were considered aviremic (<400 HIV-RNA copies/mL). Individuals H7 and H11 were reported in clinical documentation to have cognitive decline and memory impairment, raising the possibility of HAND. None of the other PDWH had a known history of neurocognitive symptoms. Five DPWH (H1, H3, H5, H6, H8) and one HIV-negative individual (N3) died with a disseminated TB infection, five showed no evidence of CNS involvement, while one (H5) was diagnosed with TB meningitis (TBM). Diagnosis was based on antemortem imaging and results: cerebellar infarction with leptomeningeal enhancement on CT brain and elevated protein and lymphocytes on CSF analysis, although the diagnosis was never microbiologically confirmed and histopathology revealed no pathological changes. Histopathological evaluation of brain core samples did not demonstrate neuropathology in any of the patients. Individuals N1 and N2 were diagnosed with a SARS-CoV-2 infection, although SARS-CoV-2 RNA was not detected in brain biopsy. In lung biopsies SARS-CoV-2 RNA was not observed for N2, by qPCR on the E and N gene of SARS-CoV-2, while for N1 spliced E gene was detectable. (**Table 1** and **Table S1**).

**Table 1.**
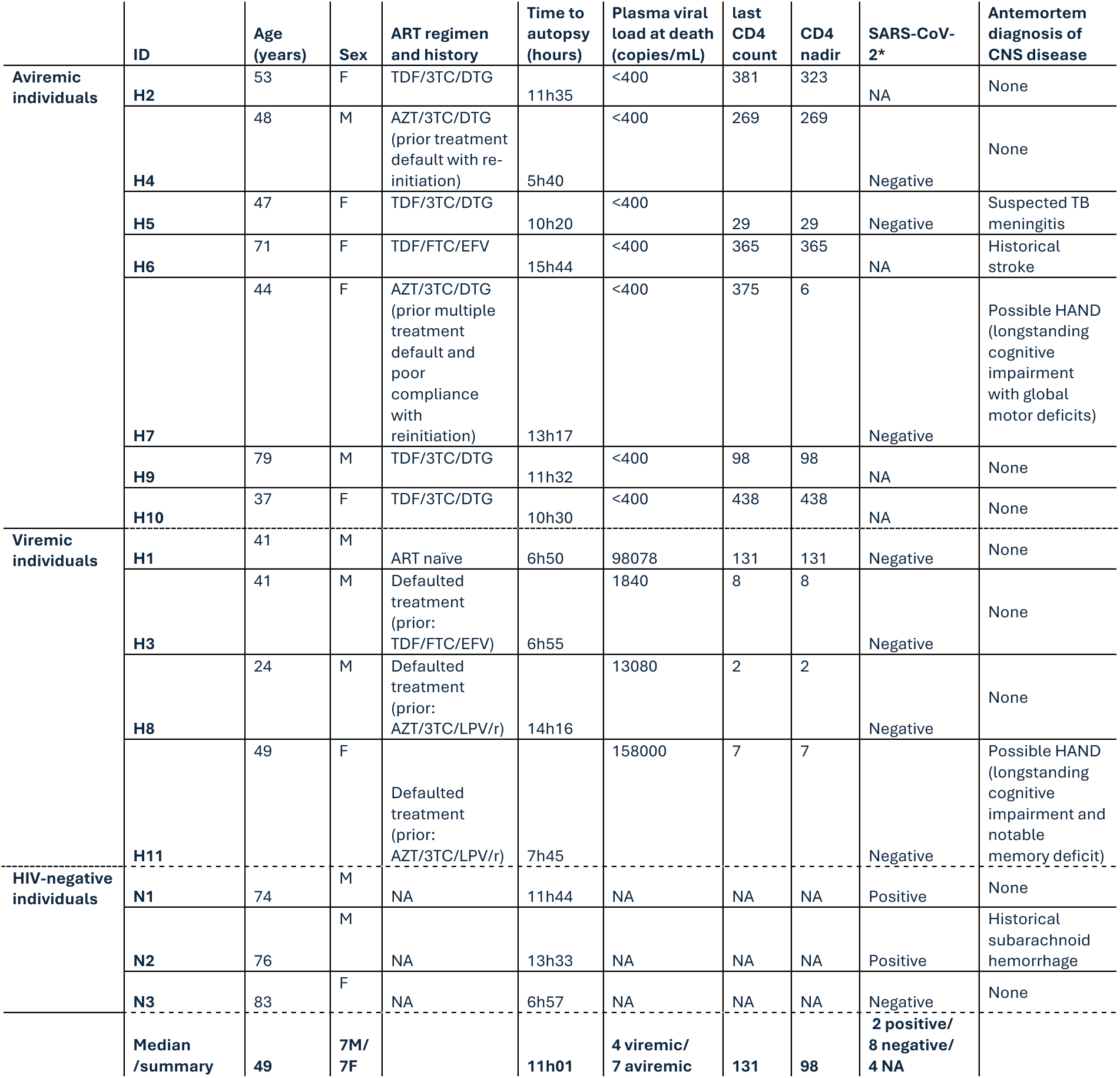
Participant characteristics. Median values or summary of the characteristics are reported in bold in the last line. The ID of individuals with HIV starts with “H”, the HIV-negative individuals start with “N”. 3TC = lamivudine, ART = antiretroviral therapy, AZT = zidovudine, CNS = central nervous system, DTG = dolutegravir, EFV = efavirenz, HAND = HIV-associated neurocognitive disorder, LPV/r = lopinavir/ritonavir, NA = not applicable, TB = tuberculosis, TDF = tenofovir disoproxil fumarate.

### Sorted nuclear fractions are cell-type specific

Using nuclei isolation and fluorescence-activated nuclei sorting (FANS), we sorted cell-type specific nuclear fractions of microglia, neurons, oligodendrocytes and the triple negative fraction, referred to as rest fraction from here onwards, of viremic (n=4), aviremic (n=7) and HIV-negative brain samples (n=3), **Figure S1**. Confocal microscopy confirmed the isolation of intact nuclei and the intranuclear staining for cell-type specific transcription factors (**Figure S2**). After nuclei sorting, we performed bulk-RNA sequencing on these nuclear fractions. Our samples had DV200 values with a median of 76%, ranging from 59%-90%, indicating high quality of nuclear RNA. Mapping resulted in 63-68% of reads being successfully mapped, in line with the presence of adapters in the raw read files, and FastQC analysis indicated consistent mapping. A median of 15,300 to 19,200 unique genes was expressed per cell type (**Table S2**).

We found distinct clustering of the samples per cellular subtype (**Figure 1A**), except for the H5-and N2-oligodendrocyte samples, which were subsequently excluded from further analyses as they were most likely contaminated with non-specific nuclear fractions (data not shown). Marker gene expression of microglia, oligodendrocyte and neurons confirmed the cell-type specificity of these sorted fractions. Additionally, the expression of astrocyte markers indicated that the rest fraction contained predominantly astrocytes (**Figure 1D**). To further confirm specificity of the sort-strategy, we conducted single-nuclei (sn)RNA sequencing. Per individual, 376 wells containing single microglia nuclei, 188 wells with single neuronal nuclei, and 188 wells with the rest fraction were sorted, oligodendrocytes were excluded. After filtering, we profiled 3,677 nuclei with a median depth of 770 reads and a median of 628 genes per nucleus. Nuclei clustered together based on sorted cell-type, further confirming that the nuclear sorting was also cell-type specific at the single-nucleus level (**Figure 1B**). Additionally, we identified 10 nuclear clusters (0-9), with 1 cluster representing neurons, 4 clusters corresponding to the rest fraction and 5 clusters corresponding to microglia, underscoring the diversity of single nuclei within these two latter fractions (**Figure 1C**). We did not observe any cell-type specific sub clustering based on HIV diagnosis or viral suppression status in our bulk-sequence data. Altogether, these results demonstrate that our nuclei isolation and sorting strategy produced high-quality data with a reliable, cell-type specific nuclei sorting approach.

**Figure 1.**
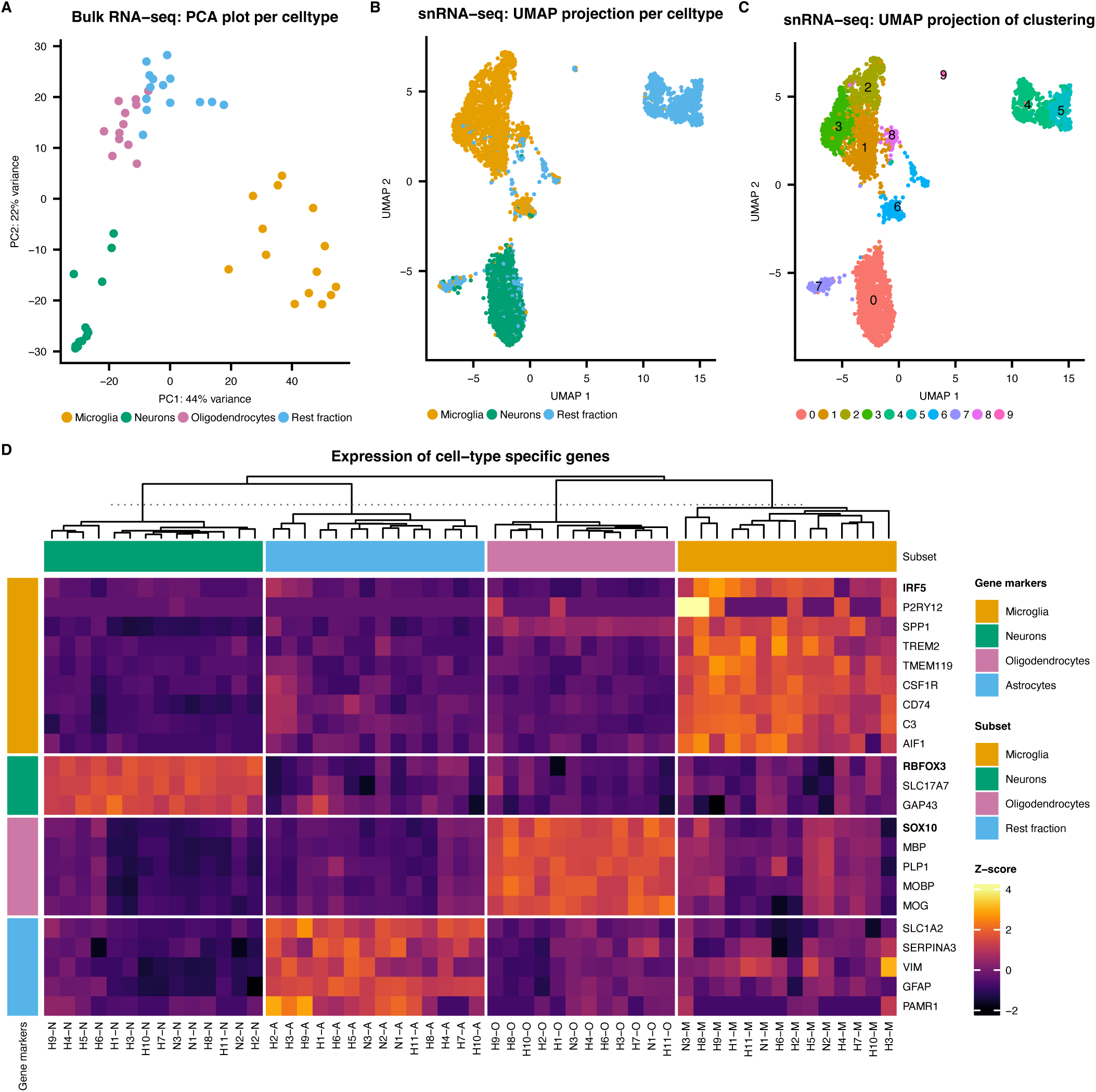
Cell-type specific of sorted data. **A)** PCA plot displaying clustering of top 500 genes of bulk-sorted samples per cell-type. **BC)** Uniform manifold approximation and projection (UMAP) plots displaying the clustering of the single-nuclei data labeled per **B)** cell type and **C)** cluster. **D)** Normalized marker gene expression of different cellular subsets. Clustering distance of columns is “Euclidean”, clustering method is “complete”. In bold text the genes that encode the transcription factors used for cell-type specific nuclear staining. (RBFOX3 is the gene coding for NeuN). Excluded samples H5-O and N2-O are not shown.

### Microglia are the predominant HIV reservoir in the brain

To investigate whether proviral DNA was present in any of the cell-type specific nuclear fractions we quantified the number of HIV LTR copies per cell-type. Proviral copies were detected in the microglial fraction of three individuals: two aviremic (H4, H9) and one viremic (H11), ranging from 1,215-10,315 copies per million cells. Trace levels of HIV were identified in microglia (H7) and in oligodendrocytes (H4) and neurons (H9). The latter two individuals also had an HIV reservoir within the microglial fraction (**Figure 2A**). Furthermore, HIV DNA was detected in the rest fraction of one additional aviremic individual (H5) (**Figure 2A**). While this fraction contains mostly astrocytes, according to our snRNA-sequencing analysis, the presence of other cell types is not ruled out. Notably, individual H5 died from TBM, which could suggest substantial T cell infiltration into the brain[39]. Analysis of CD3-epsilon expression, one of the subunits of the T cell co-receptor complex CD3, demonstrated a 4.5-fold increased expression (range: 1.2-13.5) in individual H5 compared to the other individuals, based on variance stabilizing transformation (VST) normalized and batch corrected values (**Figure 2B**). This indicates the presence of (HIV-infected) T cells in this individual. Therefore, individual H5 was removed from the data set in further analyses. In summary, we detected proviral HIV DNA in 4 aviremic and 1 viremic individual, out of 11 DPWH. Microglia served as the main cellular reservoir, although infected T cells were most likely present in individual H5. Despite screening both single- and bulk-sequencing data using both an HIV reference strain and individual specific HIV envelope sequences, derived from HIV DNA from peripheral CD4 T cells, as alignment for each individual, we could not detect HIV transcripts above background in any of the cell types.

**Figure 2.**
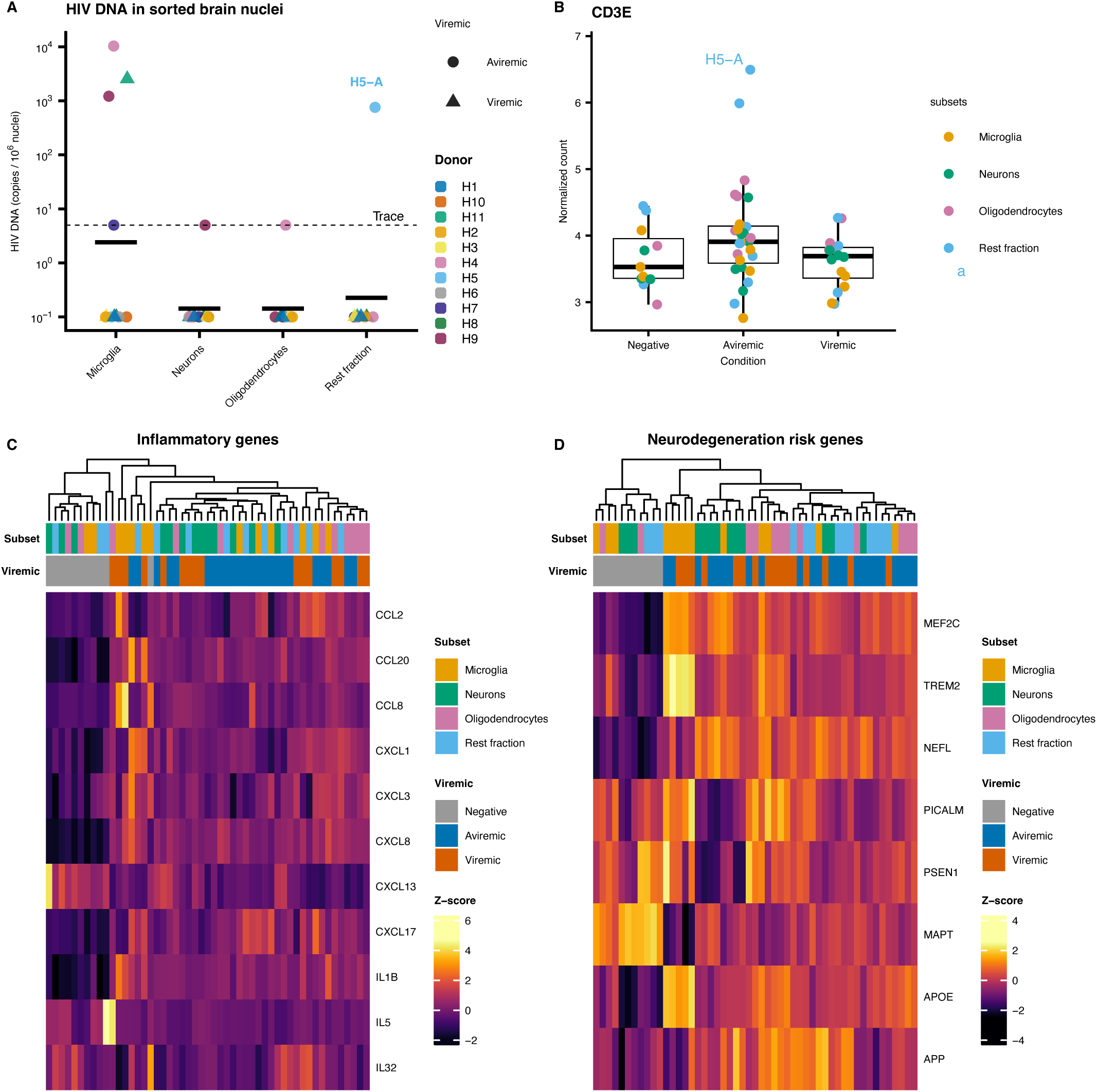
The presence of HIV reservoir by proviral LTR copies and expression of inflammatory and neurodegeneration risk genes in different cell-type derived nuclear fractions. **A)** Number of proviral LTR copies for the different cellular subsets. **B)** The expression of CD3-epsilon and **C)** Heatmap of the VST normalized and batch corrected counts of inflammatory genes. Columns are clustered by Euclidean distance. Individual H5 was excluded in the analyses for **C** and **D.. D)** Heatmap of the VST normalized and batch corrected counts of neurodegenerative risk genes. Columns are clustered by Euclidean distance. Individual H5 was not included in this analysis.

### HIV infection triggers neuroinflammation and neurodegeneration across cell types

Batch corrected and VST normalized gene counts were mapped for known neurodegeneration risk genes[40–42] and inflammatory genes previously described in *in vitro* HIV infection of microglia[32] (**Figure 2C&D**). For neurodegeneration risk genes, samples of HIV-negative individuals clustered together, demonstrating differential neurodegeneration gene expression in DPWH compared to deceased HIV-negative individuals, across all cellular compartments. The same was observed when clustering inflammatory markers, except for one microglia sample from a HIV-negative individual, which clustered within the samples of HIV-positive individuals. Both inflammatory and neurodegeneration risk genes were generally upregulated in DPWH in all cell types. This data demonstrates that in our study population HIV infection increases neuroinflammation and the risk of neurodegeneration in not just microglia, the HIV target cells, but also bystander cell-types. Furthermore, viral suppression does not seem to mitigate this increased expression, as the viremic and aviremic samples did not cluster separately.

### Elevated immune activation, cellular activity and disruption of brain development in viremic individuals compared to aviremic individuals

Suppressive ART reduces systemic inflammation that results from HIV-associated immune activation[19], though some residual inflammation persists[20–22]. Given the observed signs of neuroinflammation in DPWH, we investigated whether suppression of viral replication differentially affects neuroinflammation and function in different cell-types in the CNS. After quality control and filtering per cell-type, we profiled a total gene count of 8,198 in microglia, 12,834 in neurons, 11,382 in oligodendrocytes and in 10,261 “rest” nuclei, **Table S2.** Subsequently, we performed a differential expression analysis between HIV-negative, aviremic and viremic individuals for all 4 cellular fractions.

For viremic vs aviremic individuals, we identified 84 differentially expressed genes (DEGs) in microglia, 116 in neurons, 168 in oligodendrocytes and 183 in the rest fraction. In total, we identified 419 unique DEGs, of which 42 genes overlapped between al least 2 cell types, **Figure 3A-D** & **Table S3**. After literature review, 19 of these genes were identified as nuclear protein-coding genes and 11 were mitochondrial protein-coding genes. Most of these genes are associated with HIV (neuro)inflammation, (neuro)degeneration or immune function (**Table S4).**

**Figure 3.**
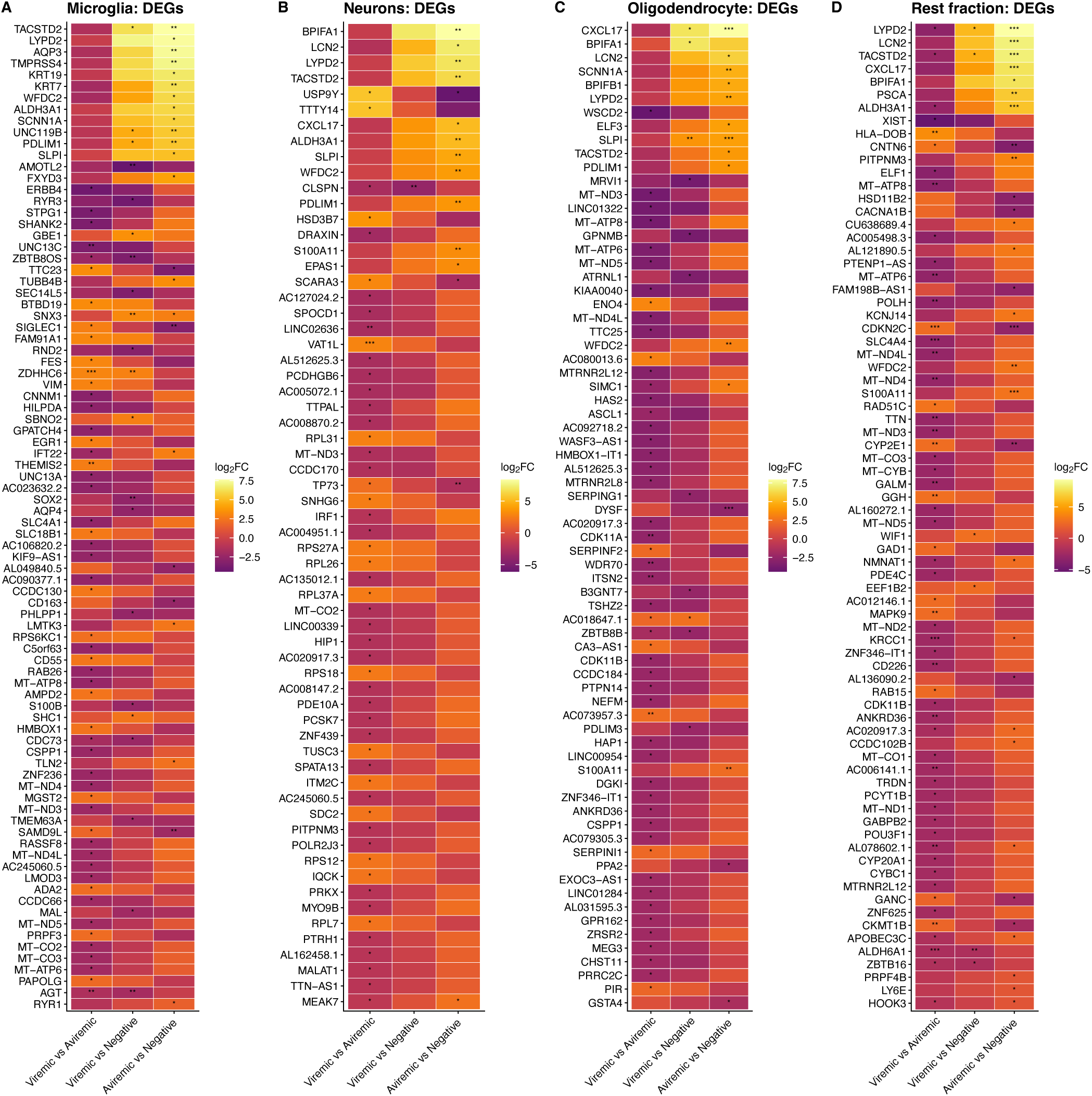
Differential gene expression per cell type-specific nuclear fraction for each comparison. **A-D:** Heatmaps for each cell-type, showing the top 50 genes per contrast with an adjusted p-value < 0.05 and LFC > 0.25 for one of the comparisons, colored by log2FC. Significant adjusted p-values are marked with asterisks (p<0.05=*, p<0.01=**, p<0.001=***).

Subsequently, a gene set enrichment analysis (GSEA) was performed per cell type, for viremic vs aviremic individuals, on the genes ranked by their LogFoldChange (LFC). An upregulation of immune response and cytokine signaling pathways was observed, specifically in the microglia of viremic individuals (**Figure 4A**). Additionally, pathways related to cytoplasmic translation were consistently upregulated across all cell types. Notably, pathways related to metabolic and biosynthetic processes were perturbated in all cell types. While GSEA showed upregulation of these pathways in neurons, the overrepresentation analysis (ORA) and DEGs showed robust downregulation of (ATP) metabolic processes and synthesis in microglia, oligodendrocytes and the rest fraction (**Figure 4A&B and Table S4)**. GSEA showed decreased expression of synaptic organization and signaling pathways in microglia and oligodendrocytes. Additionally, pathways associated with synaptic signaling and brain development were downregulated in multiple cell types (**Figure 4A&B**). Together, these findings suggest that ongoing plasma viremia in viremic individuals is associated with impaired metabolic and synaptic functions and upregulation of pathways associated with general protein synthesis and cellular activity, not limited to HIV-target cells in the brain. Additionally, in microglia increased immune responses suggest a pro-inflammatory state of these cells in viremic individuals as compared to aviremic individuals.

**Figure 4:**
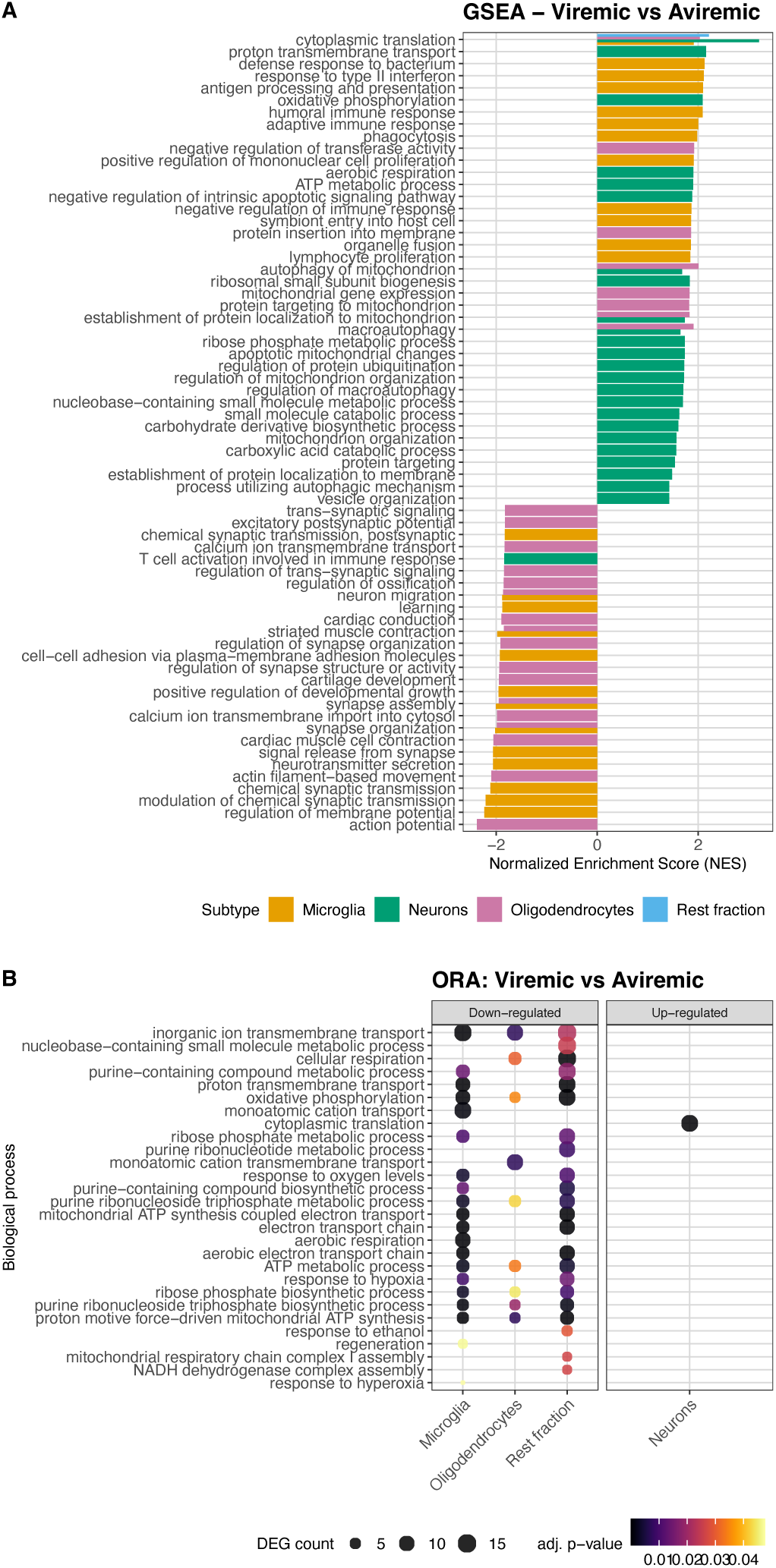
Differential expression analysis for 4 different cell types of bulk-RNA-seq data comparing viremic with aviremic DPWH. **A:** GSEA analysis with GO pathways of activated and suppressed pathways genes for all 4 cell types combined. The Normalized Enrichment Score (NES) quantifies the extent to which a specific pathway is upregulated (positive NES) or downregulated (negative NES) which is then normalized using the mean enrichment scores obtained from perputations of the same pathway, enabling to compare values between pathways. **B:** Overrepresentation analysis of up- and downregulated DEGs within GO pathways. All cell types with pathways with FDR≤0.05 included. DEG count identifies the number of DEGs in a pathway.

### Elevated immune activity across multiple cell-types compared to HIV-negative individuals

These previously discussed observations suggest that ART may protect the brain from potential brain dysfunction and increased cellular activity. However, they also raise questions about how these individuals differ from HIV-negative individuals. Within the pairwise analysis of aviremic and HIV-negative individuals, 63 unique DEGs were identified; 24 DEGs in microglia, 15 in neurons, 15 in oligodendrocytes and 31 in the rest fraction (**Figure 5A-D** & **Table S5**). When performing GSEA, an upregulation of RNA splicing pathways and a downregulation of synaptic transmission pathways was observed in mostly the oligodendrocytes and rest fraction, respectively (**Figure 5A**). Moreover, the upregulated genes were overrepresented within immune-related pathways in neurons and the oligodendrocytes (**Figure 5B**). When specifically looking to the DEGs, we found that 10 of the 11 DEGs occurring in multiple cell-types were associated with HIV (neuro)inflammation or immune function (**Table S6**).

**Figure 5.**
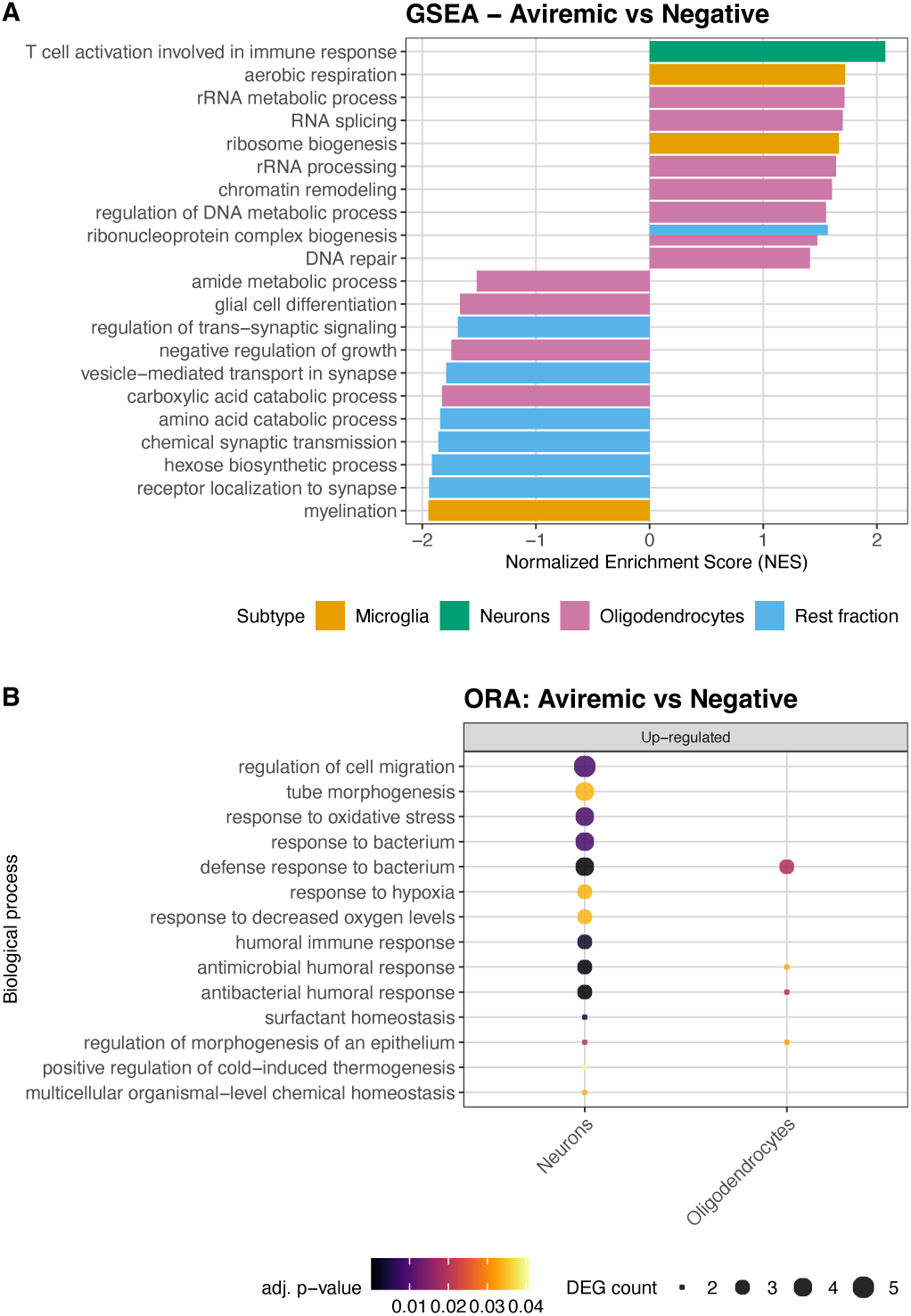
Differential expression analysis for 4 different cell-types of bulk-RNA-seq data comparing aviremic DPWH with HIV-negative individuals. **A:** GSEA analysis for microglia with GO pathways of activated and suppressed pathways genes. The NES quantifies the extent to which a specific pathway is upregulated (positive NES) or downregulated (negative NES) which is then normalized using the mean enrichment scores obtained from perputations of the same pathway, enabling to compare values between pathways. **B:** Overrepresentation analysis of up- and downregulated DEGs within GO pathways. All cell types with pathways with FDR≤ 0.05 included. DEG count identifies the number of DEGs in a pathway.

When comparing viremic DPWH with HIV-negative individuals, 21 DEGs in microglia, 1 in neurons, 11 in oligodendrocytes and 6 in the rest fraction could be observed. This resulted in 38 unique DEGs, of which 6 were identical to the DEGs in the comparison of aviremic individuals with the HIV-negative group (**Figure 6A&B** & **Table S7**). The GSEA results showed an upregulation of viral and metabolic processes in all cell types (**Figure 6A**). Additionally, an upregulation of cytoplasmic translation pathways was observed in both microglia and neurons. Moreover, synaptic signaling, growth pathways and brain development pathways were found to be downregulated, specifically in microglia. When looking specifically to the overrepresentation of the DEGs, immune related pathways were found to be upregulated in oligodendrocytes (**Figure 6B**). One of the DEGs, TACSTD2, was significant in both microglia and the rest fraction and was found to be differentially expressed in all 3 comparisons, and is known for its association with increased T cell infiltration and viral entry (**Table S6**). One other DEG, LYPD2, was found in at least one cell type in all 3 comparisons and has been associated with immune surveillance and HIV-1 susceptibility (**Table S6**).

**Figure 6.**
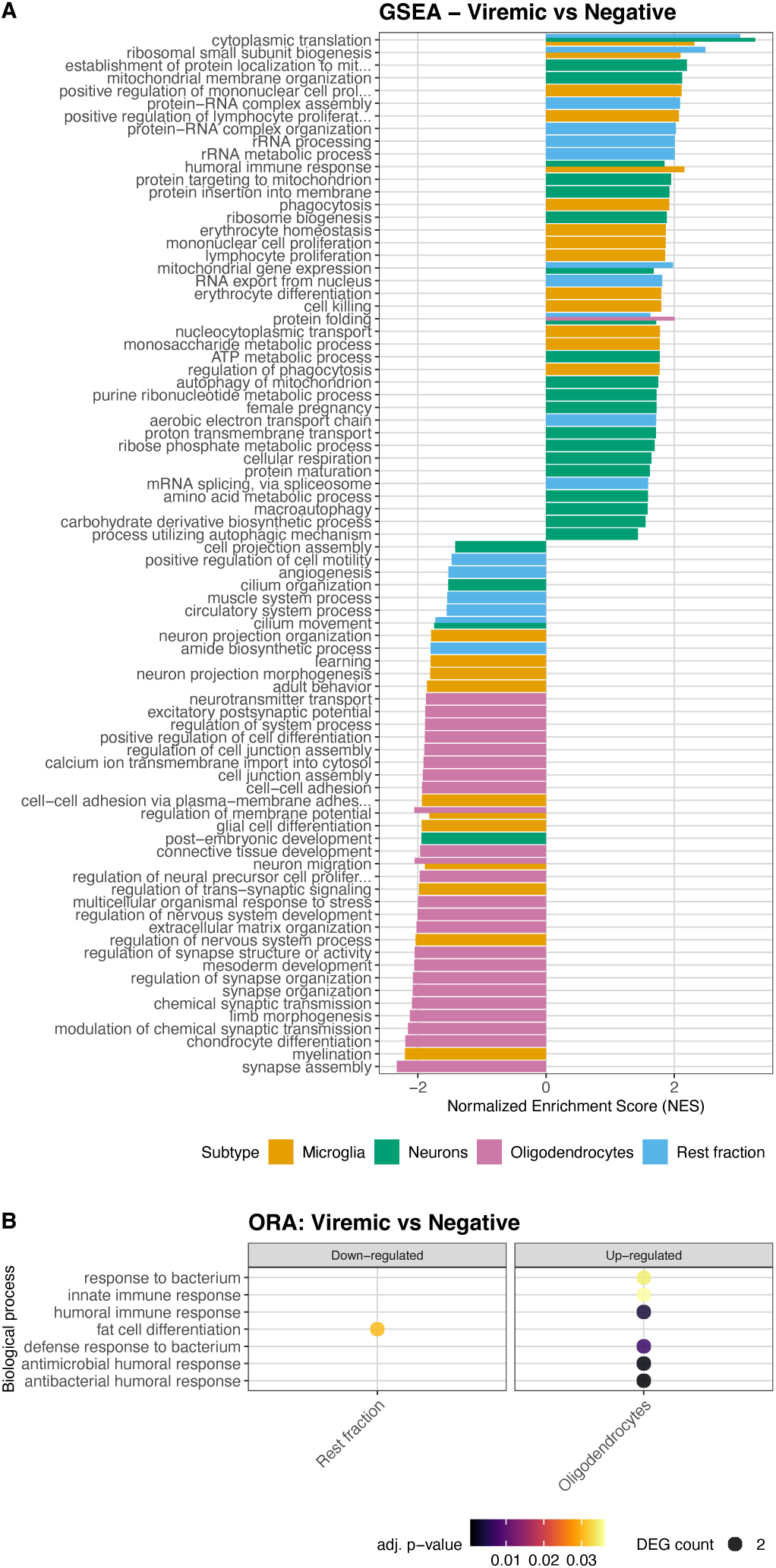
Differential expression analysis for 4 different cell types of bulk-RNA-seq data comparing viremic DPWH with HIV-negative individuals. **A:** analysis for multiple cell types with GO pathways of activated and suppressed pathways genes. The NES quantifies the extent to which a specific pathway is upregulated (positive NES) or downregulated (negative NES) which is then normalized using the mean enrichment scores obtained from perputations of the same pathway, enabling to compare values between pathways. **B:** Overrepresentation analysis of up- and downregulated DEGs within GO pathways. All cell-types with pathways with FDR≤ 0.05 are included. DEG count identifies the number of DEGs in a pathway.

These findings highlight a persistent neuroinflammatory state in both aviremic and viremic individuals compared to HIV-negative individuals. While only microglia harbor HIV-reservoirs, excluding indidual H5 with TBM, other cell types still showed increased general cellular activity and reduced synaptic functions regardless of viral suppression compared to HIV-negative individuals. Altogether, these results suggest that HIV contributes to neuroinflammation and synaptic and cellular dysfunction, and while ART was observed to act as partial protection, levels do not restore to the levels observed in HIV-negative individuals.

## Discussion

As the brain is a distinct immune-privileged anatomical sanctuary with unique resident cell types, identifying and understanding this HIV reservoir and its impact is key to advancing HIV treatment and cure strategies. In this study, we investigated the cell types in which HIV can reside and examined the cell-specific effects of HIV infection and viral suppression on neuroinflammation and brain homeostasis.

Postmortem brain material obtained shortly after death, with a median postmortem interval of 11 hours together with detailed clinical data from a well-characterized cohort[find ref] including viral load levels at time of death, provided a unique opportunity to investigate the effects of viral suppression in the brain. Using FANS, we isolated cell type-specific nuclear fractions from postmortem brain tissue of seven aviremic and four viremic DPWH, overcoming challenges typically associated with isolating these brain cells. Proviral HIV DNA was identified in microglia and is most likely also present in infiltrating T cells dependet on co-morbidities/co-infectionsm. Our results demonstrate that, while ART mitigates the effects of HIV on the brain, neuroinflammation and -degeneration persists. This remains a critical target for future HIV cure strategies and ongoing efforts to reduce morbidity and mortality among PWH despite effective ART.

We detected an HIV reservoir in five individuals (four aviremic, one viremic) out of 11 DPWH. This reservoir was predominantly located in microglia, consistent with previous findings[11, 24, 25, 43]. Nevertheless, we also identified a potential T cell reservoir in the rest fraction, in a single individual who died from TBM, which is characterized by a massive T cell influx within the brain[39]. This may imply that infected T cells may infiltrate the brain or get infected in the CNS, contributing to the persistence and dissemination of HIV within the CNS. While we did not detect an HIV reservoir in all individuals, this may be attributable to the clustered distribution of HIV within the brain, forming foci of infected cells[25], the relatively low abundance of microglia in the brain[44], limited numbers of analyzed microglial nuclei (median of 45,568 with range: 22,565-94,463), and limitations of core needle biopsy in sampling limited areas as compared to complete diagnostic autopsy[45]. Therefore it remains plausible that an HIV reservoir is present in in the CNS of all individuals, as suggested by other studies[10, 12], and that its detection strongly depends on the number of nuclei and/or the specific biopsy analyzed. Notably, viral suppression had a limited impact on the size of the viral reservoir in the brain. This is consistent with findings in all participants of the FIND2.0 study, from which the DPWH used in this study were from, where significant differences in reservoir size were only observed in lymphoid organs[46], where rapid turnover of T cells is driven by ongoing viral replication in the absence of ART [47]. Similarly, other studies have reported comparable levels of viral reservoirs in the brains of both aviremic and viremic individuals[10, 11]. Unlike CD4^+^ T cells in peripheral blood, microglia are more resistant to cell death[48], have longer lifespans[49] and are subject to reduced immune-mediated cell killing and diminished immune surveillance within the CNS[50]. These low turnover rates exhibited by microglia could render them less dynamic and more persistent in the presence of ART than (peripheral) T cells. This persistence may have critical implications for PWH on ART and for the development of future HIV cure strategies.

The observed transcriptomic changes and the persistence of HIV reservoirs in the brain suggest the presence of ongoing viral transcription and production, which may drive neuroinflammation and contribute to the development of HAND[7–9]. Likewise, recent studies in DPWH reported a downregulation of homeostatic and metabolic pathways in microglia and astrocytes, compared to the HIV-negative individuals[35, 36], underscoring the detrimental effect of neuroinflammation on brain function. However, the specific impact of ART on neuroinflammation and brain homeostasis was still unknown. Previous studies have shown that while ART effectively suppresses ongoing viral replication and systemic inflammation[19], residual viral transcription and production persists, driving inflammation[20–22]. In line with these findings, our results demonstrate that viral suppression by ART is associated with downregulation of pathways related to homeostasis and synaptic signaling across multiple brain cell types. Additionally, we identified an upregulation of immune-related pathways and general cellular activity across brain cell types. These patterns were also observed in both aviremic and viremic individuals when compared to HIV-negative individuals. Altogether, these results suggest that HIV infection induces neuroinflammation with pronounced effects in microglia, the primary target cells, impacting not only the infected cells but also neighboring bystander cells involved in maintaining healthy brain function. Although viral suppression by ART mitigates these effects, pathways are not fully restored, possibly caused by persistent low-level viral activity during treatment. This may explain why HAND persists in individuals on ART, albeit with reduced severity[6]. While our findings are suggestive for impaired brain function in DPWH based on transcriptomic data, we lacked sufficient clinical evidence to establish causality with the current sample size and clinical data. Future studies with larger cohorts and direct clinical assessments are needed to validate these findings and elucidate the relationship between (residual) HIV transcription, neuroinflammation, viral suppression and HAND.

In our study, viral suppression was defined peripherally in plasma, and no viral transcripts were detected within the brain. Two recent studies investigating single nuclei of DPWH detected HIV transcripts only in microglial of HIVE individuals[35, 36]. Future analyses using specialized amplification and sequencing approaches targeting HIV transcripts, and sequencing of additional brain material are necessary to conclusively determine the presence of viral transcripts.

The reported presence of an HIV reservoir within the brain suggests potential local CNS effects of viral transcription and production on neuroinflammation. Immune pathways were upregulated in all cell types, arguing for a systemic brain effect on neuroinflammation. This effect was more pronounced in microglia, the predominant cellular reservoir for HIV in the brain, in viremic relative to aviremic individuals. However, when comparing the brains of those with detectable HIV DNA and those without, no significant differences in pathways associated with neuroinflammation or brain functioning were observed. This may be attributed to the low number of nuclei analyzed and the focal distribution of HIV infection in the brain[25], as previously discussed. All individuals might have a reservoir, although confined to specific areas, while the effect of HIV infection on neuroinflammation appears to be more diffused throughout the brain. Although we did not distinguish between intact and defective proviruses by assessing total HIV DNA, defective viruses have been shown generate viral transcripts and viral protein thereby inducing immune activation[22, 51], thus their specific effects should be considered too. In conclusion, our findings suggest that neuroinflammation may result from a combination of local and systemic HIV transcription. Future research focusing on distinguishing between these local and systemic contributions to neuroinflammation may clarify the mechanisms driving neuroinflammation in both aviremic and viremic PWH.

The use of FANS provides unique opportunities to study cellular reservoirs from postmortem brain tissue, with nuclear transcriptomic profiles serving as a good reflection of cellular transcriptomic profiles[52, 53]. However, challenges remain, including differences in RNA distribution between the cytosol and the nucleus[52, 54] and limited markers to isolate diverse cell types. Beyond microglia, other potential resident cellular reservoirs in the brain have been described, including perivascular macrophages[25], pericytes[55] and astrocytes[24, 43]. IIn our study, microglia were sorted based on IRF5 expression, a marker also present on other myeloid cells. Consequently, it remains uncertain whether the observed HIV reservoir within microglia also includes perivascular macrophages. Furthermore, our transcriptomic data revealed that astrocytes, pericytes but also T cells were sorted in the rest fraction, preventing their specific isolation and analysis for cell-type specific effects. Future studies should focus on identifying nuclear markers to better isolate and study these potential HIV reservoirs in the brain. Additionally, more in-depth single-nuclei studies are needed to address the substantial heterogeneity of microglia and other cellular reservoirs in the brain[56]. For example, clusters of activated microglia, known as disease-associated microglia (DAM), which are linked to neurodegenerative diseases, have been identified[57], as well as clusters of reactive astrocytes in HIV infected brains[36]. Investigating whether HIV infection is linked to these specific populations of cells would provide valuable insights into its role in neuroinflammation and brain pathology.

Finally, our study cohort size posed limitations. Previous literature indicates that transcriptomic data in human brain studies are often influenced more by technical and host-related factors, such as age and sex, than by diagnosis[56]. A proportion of our participants were diagnosed with TB, and some HIV-negative individuals were infected with SARS-CoV-2. Specifically, for the individual with TBM, it is known that associated neuroinflammation and the microglial involvement in the response to this infection might impact the transcriptomic data[58], which is why we excluded samples of this individual from the downstream analysis. The small sample sizes (HIV-negative; n=3, viremic; n=4, aviremic; n=6) may have contributed to the under or overestimation of gene and GO-term significance. Additionally, the HIV-negative individuals were significantly older than the DPWH, potentially exhibiting higher levels of chronic immune activation, termed inflammaging[59]. Despite this, we still observed upregulation of immune-related pathways in DPWH compared to the HIV-negative individuals, suggesting a robust and extensive shift in transcriptome. Furthermore, baseline immune and cellular pathway upregulation in the HIV-negative group may have reduced detectable DEGs versus DPWH, while similar ages among viremic and aviremic DPWH limit these inflammaging effects, yielding a larger DEG set in that comparison. Moreover, the HIV-negative group was also composed of entirely male participants, which may have influenced levels of immune response and inflammation as well[60, 61]. Despite these potential confounders, we did not find significant clustering differences based on TB, SARS-CoV-2, sex or age in our analysis based on PCA plots. The identification of significant results related to HIV infection status within our cohort is promising, and future studies with larger cohorts could validate and expand upon our findings.

Though HIV persistence in the brain during ART has been recognized for years, the question of how viral suppression shapes HIV persistence and residual neuropathology has remained unanswered. Our study addresses this gap directly, being, to our knowledge, the first to define the impact of viral suppression on the CNS HIV reservoir and transcriptional state of various brain cells by comparing postmortem brain tissue of viremic and aviremic DPWH. By the use of FANS, we uniquely dissected the cell-type specific transcriptional patterns between viremic states in the main cell-types of interest (microglia, neurons, oligodendrocytes and rest fraction). Our findings highlight the presence of the proviral HIV reservoir in microglia and the impact of HIV infection on these target cells and bystander cell-types in the CNS, despite ART-induced viral suppression. While viral supression reduces the effects of neuroinflammation and synaptic and metabolic disruption in both HIV target cells and other brain cells, thereby potentially mitigating effects of HIV on brain health, pre-disease homeostasis in the brain is not achieved. These insights emphasize the need to consider the impact of HIV, ART and viral transcription on the brain when developing treatment and cure strategies. Moreover, they underscore the importance of monitoring the CNS to ensure the effectiveness and safety of these interventions, especially HIV cure interventions that stimulate viral transcription.

## Acknowledgements

We gratefully acknowledge the contribution of Justine Blonk to this work. Justine was involved in the development of the study protocol and in data collection, and sadly passed away before the completion of this manuscript. We wish to honour her contribution and express our sincere appreciation for her efforts. We acknowledge the Utrecht Sequencing Facility (USEQ) for providing sequencing service and data. USEQ is subsidized by the University Medical Center Utrecht and The Netherlands X-omics Initiative (NWO project 184.034.019). Single Cell Discoveries provided helpful advice regarding library preparation, sequencing alignment and data analysis. We acknowledge Alistair Calver and Gajendra Chita for serving on the panel for clinicopathological conferences and thoroughly investigating participants’ clinical data. We acknowledge the North West Provincial Department of Health for approving our autopsy study, and the Klerksdorp Tshepong Hospital Complex for allowing and facilitating sample collections. We would also like to thank the lab of Dr. Dracheva at the James J, Peters VA Medical Center, Bronx, NY, 10468, US for advice and support on the nuclei isolation and FANS technique. We acknowledge Weizhen Xu (UMC Utrecht) for his help with sequence alignment, Dani Heesterbeek, Remy Muts and Marije van ‘t Wout for their help with confocal microscopy, and the Core Flow cytometry Facility (CFF) for their help with flow cytometry. We thank all the participating individuals for their generous and selfless contribution to this study. Lastly, we thank the family members of deceased patients for providing consent for us to postmortem sample their loved ones.

## Author contributions

JS, MN and LW designed the study. NS, AB, EV, WDFV, NM, AW, MP and MN performed and organized sample collection. TV performed histopathologic analyses. MMN, NZ, MM, SG, PS and CMS performed experiments. MMN, NZ and KA performed analyses under the supervision of RK, LW and JS. MMN and NZ wrote the manuscript and all authors revised and approved it.

## Data availability

Data available upon reasonable request.

## Methods

### Human tissue

Deceased human individuals were sampled within the framework of a larger study – the FIND2.0 study: Core needle biopsies to detect the hidden HIV reservoirs in hard-to-reach tissue compartments of ART-suppressed and uncontrolled viremic PWH[62]. The FIND2.0 study comprised rapid post-mortem minimally invasive tissue sampling of viremic and aviremic DPWH, after obtaining qualified next-of-kin informed consent for specimen collection. The objective of FIND was to perform in-depth characterization of the HIV reservoir in multiple organ tissues and fluid compartments. The study used minimally invasive autopsies with fine-needle and core biopsies on individuals above 18 years of age who died while hospitalized at the Klerksdorp-Tshepong Hospital, North-West Province, South Africa[62].

This study was performed on human autopsy brain samples from 11 FIND2.0 study participants for whom sufficient brain material was available. In addition, brain samples from 3 deceased HIV-negative individuals from a parallel study of patients with presumptive pneumonia who died in hospital[63] in the same population were used as HIV-negative controls, **Figure S1 and Table 1**. These HIV-negative individuals were sampled by the same physician at the same research center and using similar methodology and patient selection criteria[63]. Autopsies were performed within 16 hours of death, with a median of 11h01 (IQR 6h57-13h17). HIV status was defined by documentation of an HIV ELISA or HIV point-of-care rapid test (if positive repeated with a confirmatory second rapid test according to local guidelines)[64]. For those receiving ART (n=7), a documentation of receipt of therapy and a viral load measurement either prior to or at autopsy was required, whereas those not on ART and HIV positive (n=4) were required to either be ART-naïve or have no documented ART exposure for >6 months prior to death. Individuals with plasma viral load of <400 copies/mL at autopsy, along with evidence of ART adherence were considered virally suppressed (aviremic). HIV-negative individuals were included during the COVID-19 pandemic as controls; they were required to be negative for SARS-CoV-2 envelope (E) and nucleocapsid (N) RNA in their brain biopsies. Individuals who otherwise would fulfill eligibility criteria, but who had confirmed premortem diagnoses of hematological malignancies or intracranial infections/lesions were excluded to minimize confounding from pronounced immune activation or infiltration, however individuals who had a suspected but unconfirmed diagnosis were still eligible for inclusion. Detailed demographic and clinical data were collected from admission and consolidated during clinicopathological conferences (described below), found in

**Table 1** and **Table S1**.

In the presence of an HIV counsellor, the legally determinded next-of-kin provided written informed consent postmortem ahead of study procedures[62]. Consent was obtained by a medical doctor and a trained HIV counsellor in the preferred language of the relative. Ethical approval for the FIND study and the PotPrev study was obtained from the University of the Witwatersrand Human Research Ethics Committee (reference numbers 161113 and 171009 respectively) as well as by both provincial and hospital research committees.

### Postmortem sampling and clinical review

Postmortem sampling was performed by a single study physician. Brain biopsies were taken via core needle biopsy within 16 hours of death. Biopsy procedures used an aseptic procedure. A co-axial needle/introducer was inserted into the brain via a trans-orbital approach using a 16-gauge core biopsy needle and the BARD Magnum reusable biopsy instrument (BARD, Covington, USA). This procedure was performed bilaterally with 5 cores obtained from each hemisphere while advancing the depth of the needle to ensure collection of tissue from several areas. This tissue was immediately preserved in cryotubes in an ethanol dry ice-slurry until it could be frozen at - 80°C (within approximately 2 hours). Subsequently, it was transported on dry ice to the Netherlands and stored again at -80°C until further use. A portion of each core biopsy, sectioned ahead of storage at -80°C, was fixed in 10% formalin and submitted for histopathological review by a senior academic histopathologist with experience in postmortem samples to confirm both tissue type and pathology present. SARS-CoV2 PCR testing was performed on nasopharyngeal swabs obtained during the participant’s admission to hospital or postmortem where not available.

Concurrently, a review of participant primary care clinic cards (where available) and hospital records, electrocardiography, echocardiography, radiological (x-rays, CT scans, MRI, ultrasonography) and laboratory investigations (blood results, microbiology and histology reports), **Table S1**, was undertaken by three senior academic specialist physicians independently to determine pathologies present and immediate, underlying and contributory causes of death. Over the course of the study, clinicopathological conferences were conducted in which consultation of a panel consisting of these physicians and histopathologist together with all supporting material was undertaken and immediate, underlying and contributory causes of death as per World Health Organisation guidelines documented[65]. Detailed results are described elsewhere[62]. Particular emphasis was placed on diagnosis of tuberculosis which is endemic in this population and may be clinically diagnosed and treated without microbiological confirmation[66, 67].

### Nuclei isolation and sorting

Nuclei were isolated from each individual’s frozen brain biopsies, randomly selecting six biopsy cores (maximum of 500 mg) per individual as previously described with some adjustments[35, 68], **Figure S1**. Tissue was homogenized by douncing using a glass Dounce homogenizer (D9063, Sigma-Aldrich) in ice-cold lysis buffer (0.32M Sucrose (Sigma Aldrich, S0389), 5mM CaCl_2_ (Sigma-Aldrich, 2115), 3mM Mg(CH_3_COO)_2_ (Sigma-Aldrich, 63052), 0.1mM EDTA pH8.0 (Invitrogen, AM9260G), 10mM Tris-HCL pH8.0 (Invitrogen, 15568025), 0.1% Triton X-100 (Sigma-Aldrich, 93443), 1mM DTT (Sigma-Aldrich, 43816), 400U/mL RNAse inhibitor (Takara, 2313A) in nuclease-free water). The homogenate was filtered using 40μm strainers (732-2757, VWR), layered under ice-cold sucrose buffer (1.8M Sucrose, 3mM Mg(CH_3_COO)_2_, 10mM Tris-HCL pH 8.0, 1mM DTT, 400U/mL RNAse inhibitor in nuclease-free water) and ultracentrifuged at 70,000 RCF for 1 hour at 4°C (Sorvall, MTX 150 Micro-Ultracentrifuge, Thermo Fisher). Pelleted nuclei were recovered in ice-cold blocking buffer (1% BSA (Sigma-Aldrich, A4503), 400U/mL RNAse inhibitor in PBS).

### Flow cytometry

Nuclei were stained (NeuN-PE, Clone A60, Sigma-Aldrich; SOX10-AF647, clone SP267, Abcam; IRF5-AF488, #IC4508G, Bio-Techne) for 1 hour at 4°C. Shortly before acquiring nuclei on the flow cytometer, 2.8uM DAPI (MBD0015, Sigma Aldrich) was added to stain all nuclei. Nuclei sorting was performed on a MA900 Cell Sorter (SONY) using a 70μm nozzle and a pre-cooled sample and collection chamber, **Figure S1**. Cell-type specific nuclei of different fractions were sorted: neurons (DAPI^+^, NeuN^+^), oligodendrocytes (DAPI^+^, NeuN^−^, SOX10^+^, IRF5^−^), microglia (DAPI^+^, NeuN^−^ , SOX10^−^, IRF5^−^) and a triple negative “rest” fraction (DAPI^+^, NeuN^−^, SOX10^−^, IRF5^−^)[68]. Bulk-sorted nuclei were sorted into 750μl ice-cold buffer (2% BSA + 400U/mL RRI in PBS), with a maximum of a 1:1 dilution of the buffer (approximately 300,000 nuclei per tube). Single-sorted nuclei were sorted into ice-cold 384 well-plates provided by Single Cell Discoveries, enabling later SORT-seq (described below). Bulk-sorted nuclei were centrifuged at 6000g for 15 minutes at 4°C and resuspended in the correct buffer depending on the downstream analysis.

### Nucleic acid isolation

DNA was extracted from the bulk-sorted nuclei using the Monarch Genomic DNA Purification Kit (New England Biolabs) according to manufacturer’s instructions. RNA was extracted from bulk-sorted nuclei via the TRIzol Reagent (15596026, Invitrogen) with only 1/5^th^ of the volumes as described by the manufacturer’s instructions because of expected low RNA yields. GlycoBlue Coprecipitant (AM9516, Thermo Fisher) was used to visualize RNA pellets.

### HIV-1 LTR quantification

The size of the proviral DNA reservoir was determined by quantifying the number of HIV-1 LTR copies via the ultrasensitive quantification droplet digital PCR (ddPCR) method as described before[69, 70]. Samples with 0-1 positive droplets were classified as negative, those with 2-4 positive droplets were classified as containing “traces” of proviral DNA, and samples with 5 or more positive droplets were considered positive for proviral DNA. Analysis of the results was performed with Quantasoft™ (BioRad), for which replicate wells were combined before analysis. Graphs were generated with GraphPad Prism (Dotmatics).

### RNA sequencing

Bulk-sorted samples were divided over 2 libraries, depending on their RNA concentrations. Single-nuclei sequencing was performed according to the SORT-seq method[71]. Each SORT-seq plate generated one library. Libraries were generated by Single Cell Discoveries (Utrecht, The Netherlands) based on the CEL-seq2 method. This enabled early barcoding, highly multiplexed analysis and 3’ end tagging to enable accurate expression levels without having to account for gene length and requiring lower sequencing reads (10M/bulk sample, 28M per plate)[72]. Quantity and quality of libraries were analyzed with the Bioanalyzer (Agilent), among which the DV200 values. The DV200 value represents the percentage of reads that are over 200 nucleotide in length, with samples having DV200 values above 30% considered to have good transcription quality[73]. Libraries were paired-end sequenced (2x 50bp) on a NextSeq2000 flow cell (Illumina) at Utrecht Sequencing Facility (USEQ).

### Data processing

#### Bulk-seq

Sequencing alignment was performed by Single-Cell Discoveries (Utrecht, The Netherlands). The raw sequencing data per library (FASTQ files) were mapped against the human genome (GRCh38, ENSEMBL release 99) and, independently, to a custom HIV-specific reference genome using STARsolo (version 2.7.11b). Mapped reads towards their genes were counted to generate the count matrices using an in-house R script that replaced the sample barcode with the sample name. Mapping against the human genome allowed up to one mismatch. Library FASTQ files were demultiplexed per sample using the FQTK (version 0.3.0), resulting in FASTQ files per sample. During this process, the R1 information, including the sample barcode, was moved to the header of R2, resulting in artificial single-end data.

#### SORT-seq

Raw SORT-sequencing data were preprocessed by removing the Illumina adapter sequences and homopolymers. Subsequently, library alignment was mapped against the human genome (GRCh38, ENSEMBL release 99), and, independently to a custom HIV-specific reference genome using STARsolo (version 2.7.11b). Cell barcodes (CB) were matched against the CB whitelist, and barcode correction was applied if needed, allowing for up to one mismatch when aligning to the human genome. CB demultiplexing and Unique Molecular Identifier (UMI) deduplication were performed, and the deduplicated UMIs were counted towards their gene/cells, generating count matrices.

### HIV RNA detection

The HIV reference was constructed using the complete HXB2 reference sequence (NC_001802.1), including HIV-Tat, HIV-LTR, HIV Tat and Long LTR transcripts[74] adapted to subtype C, as well as specific envelope sequences obtained from blood of the participant. BLAST+[75] was used to align these sequences to the human genome GRCh38, and any transcripts that showed alignment were excluded from the collection. Multiple mapping reads were also included within this analysis. The sequences that did not map to the human genome were then aligned to the HIV viral reference strains HXB2, HIV subtype C and participant specific envelope sequence. As a control the sequences from the HIV-negative individuals were also used to align against HIV strains and envelope specific sequences. The number of positive alignments in the negative control were set as the background.

### Data analysis

Analyses were performed using R (v4.5.2, 2025-10-31[76]) and Rstudio(v2025.9.2.418[77]). Plots were generated using ggplot2(v4.0.1[78]) and ComplexHeatmap(v2.26.0[79, 80]) and formatted using cowplot(v1.2.0[81]). Colors from viridisLite(v.0.4.2[82]) were used for colorblind safe figures. All analyses involving stochastic components (including enrichment analyses) were performed with fixed random seeds (set.seed in R) to ensure reproducibility.

#### Bulk-RNA-sequencing

Raw gene-level count matrices and curated sample metadata were imported into DESeq2(v1.50.2[83]) and used to construct DESeqDataSet objects via DESeqDataSetFromMatrix. For viremic-status analyses, a design formula of ∼ viremic + library was used. As standard part of and according to the internal DEseq2 workflow, size factors were estimated using the median-ratio method, dispersions were estimated by empirical Bayes shrinkage, and negative binomial generalized linear models were fit. A filtering criterium was applied to the included genes within each subset to reduce the risk of low-level contamination and false-positive results. For microglia, a minimum expression of 10 was required in half of the samples, whereas this was 20 for other cell-types because of higher RNA input and higher numbers of unique genes (**Table S1**). Wald tests were used to assess differential expression for the following contrasts: Viremic vs. Negative, Aviremic vs. Negative, and Viremic vs. Aviremic. P values were adjusted using the Benjamini–Hochberg (BH) false discovery rate (FDR) procedure[84], and log2 fold changes were stabilized using empirical Bayes shrinkage with a normal prior (lfcShrink(type = “normal”)). Analyses were performed independently for each cell type.

To prepare expression count data for visualization, normalized counts were transformed using the variance stabilizing transformation (VST)[85]. The transformation was applied after size-factor estimation. Two samples suspected of contamination with nuclei from other fractions (H5-O and N2-O) were removed prior to transformation to maintain consistency across analyses. Principle component analysis (PCA) was conducted to assess possible confounder effect of our metadata on the expression counts. Sample clustering had signs of clustering per RNA library which was therefore included in differently expressed analysis as confounding factor. Tuberculosis (TB) or SARS-CoV-2 infection, day of the performed experiment, age and sex were not found to have separate clustering and were therefore not considered as confounding factor. Batch correction was performed using removeBatchEffect from limma(v3.66.0[86]). Library preparation batch was included as the primary batch variable, while additional biological and technical factors (subsets, donor, viremic status, tuberculosis status, DNA yield, reservoir size, HIV status, and isolation day) were modeled as covariates: Y_corrected_ = Y_VST_ − batch effects − covariate effects. A design matrix modeling viremic status (∼ viremic) ensured that the correction procedure did not remove the biological signal of interest.

The VST normalized and batch corrected counts were used for generation of the cell-type dependent PCA plot, and primary data exploration (CD3E count plot and heatmaps for inflammatory and neurodegeneration risk genes). Gene expression patterns were explored with a list of genes of interest. These included well-known HIV markers, as well as markers at risk for neurodegeneration selected based on Alzheimer’s Disease studies:“TREM2”, “APOE”, “APP”, “MAPT”, “NEFL”, “MEF2C”, “PICALM”, “PSEN1”[40–42], and inflammatory genes previously described in *in vitro* HIV microglial cultures: CCL2/8/20, CXCL3/8/13/17 and IL1B/5/32[32]. Individual H5 had died from tuberculosis meningitis (TBM), and primary exploration of CD3E counts after VST normalization and batch correction suggested infiltration of T cells in the brain. The extensive CNS perturbation in this individual was reason to remove this individual from the dataset for further analysis.

Differentially expressed genes (DEGs), genes passing differential expression criteria (adjusted p-value ≤ 0.05, (absolute) LFC >0), were extracted from the DESeq2 object. Literature research was performed via Pubmed.gov by checking the DEGs in combination with the search terms “HIV”, “inflammation”, “infection”. If no results were positive, the last two were replaced by “CNS”. Subsequently, differential expression analyses were performed per sorted cell-type, comparing aviremic, viremic and HIV-negative indiviuals. Firstly, gene-set enrichtment analysis (GSEA) was performed using clusterProfiler(v4.18.2[87]) with the gseGO function on preranked gene lists derived from DESeq2 shrunken log2 fold changes. Gene symbols were mapped to Entrez Gene IDs using org.Hs.eg.db(v3.22.0[88]). Analyses were conducted against the Gene Ontology (GO) Biological Process (BP) ontology, using a weighted Kolmogorov–Smirnov–like enrichment statistic. Gene sets were restricted to 50–500 genes. No internal significance cutoff was applied (pvalueCutoff = 1), allowing all pathways to be tested. Post hoc significance was determined using BH FDR correction, retaining pathways with FDR ≤ 0.05. Redundant GO terms were simplified based on semantic similarity (cutoff = 0.8), retaining the most statistically significant representative term.

Next, overrepresentation analysis (ORA) was performed, using clusterProfiler, separately for up-and down-regulated genes using a post hoc filtering approach. DEGs were tested against the set of all genes passing DESeq2 filtering for the corresponding cell type. Again, gene symbols were mapped to Entrez Gene IDs using org.Hs.eg.db. Analyses were restricted to GO BP terms with 10–500 genes. ORA was performed without internal p-value filtering (pvalueCutoff = 1; qvalueCutoff = 1), and significance was determined post hoc using FDR ≤ 0.05. Redundant pathways were reduced using semantic similarity–based simplification (cutoff = 0.8), and pathways supported by fewer than two DEGs were excluded. ORA results were visualized as separate up- and down-regulated pathway sets, with top-ranked terms per contrast and cell type selected for display.

#### SORT-seq

Single nuclei count matrices were filtered to eliminate cell barcodes containing ambient RNA (empty wells) and filter nuclei with low-quality or high contamination based on percentage mitochondrial DNA (<5), UMI count (>350), counts of spike-in RNA molecules (>200), and the number of genes (>350) detected in each nucleus/well. Seurat(v5.4.0[89]) was used to perform normalization and scaling of the data[90, 91]. Uniform manifold approximation and projection (UMAP) dimensional reduction and graph-based clustering were performed with the first 15 principal components and a resolution of 0.5 on the scaled data, using Seurat and SeuratObject(v5.3.0[92]).

### Confocal microscopy

Frozen postmortem control brain tissue was provided by the Netherlands Brain Bank (NBB) of one individual (protocol 1233S2) for visualization of the nuclear fluorescent staining, to confirm intact nuclei upon isolation. This individual provided informed consent before participation in the study. Nuclei were stained (NeuN-AF488, Clone A60, Sigma-Aldrich; SOX10, clone SP267, Abcam; IRF5-AF488, #IC4508G, Bio-Techne; mCLING-AF488&647 clones 710 006AT488 & 710 006AT647N, SYSY Antibodies) for 1 hour at 4°C. Subsequently, nuclei were centrifuged at 6000g for 15 minutes at 4°C, resuspended in PBS and positioned on pre-treated Poly-L-lysin (P4707, Sigma-Aldrich) coverslips (543079, Greiner). Images were obtained using a Leica SP5 confocal microscope using a HCX PL APO 63x/1.40-0.60 OIL objective (Leica Microsystems).

## Supplemental Figures

**Figure S1.**
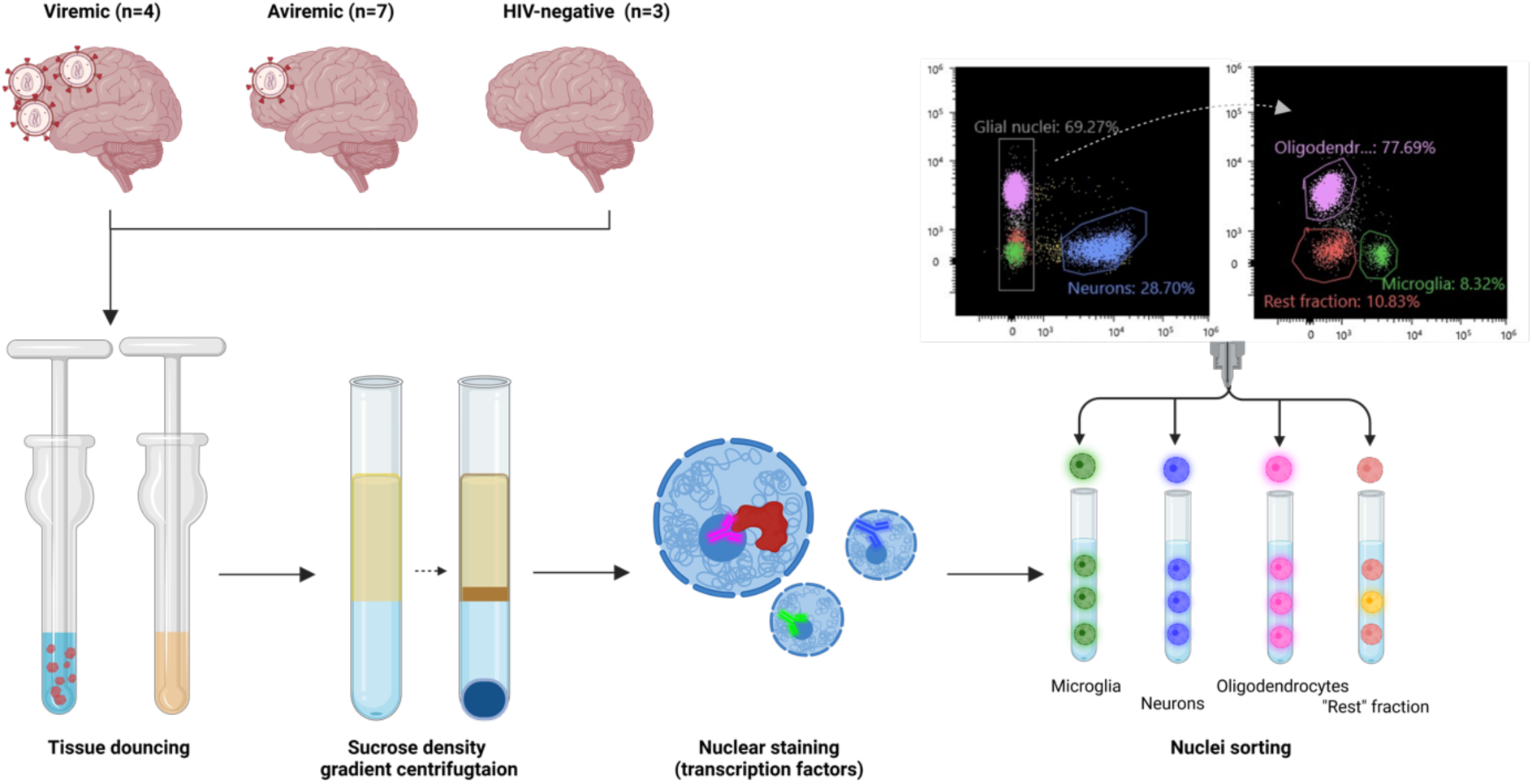
Workflow of nuclei isolation and sorting. Representative FANS plot for sorted nuclear fractions are shown. **Created with** Biorender.com

**Figure S2.**
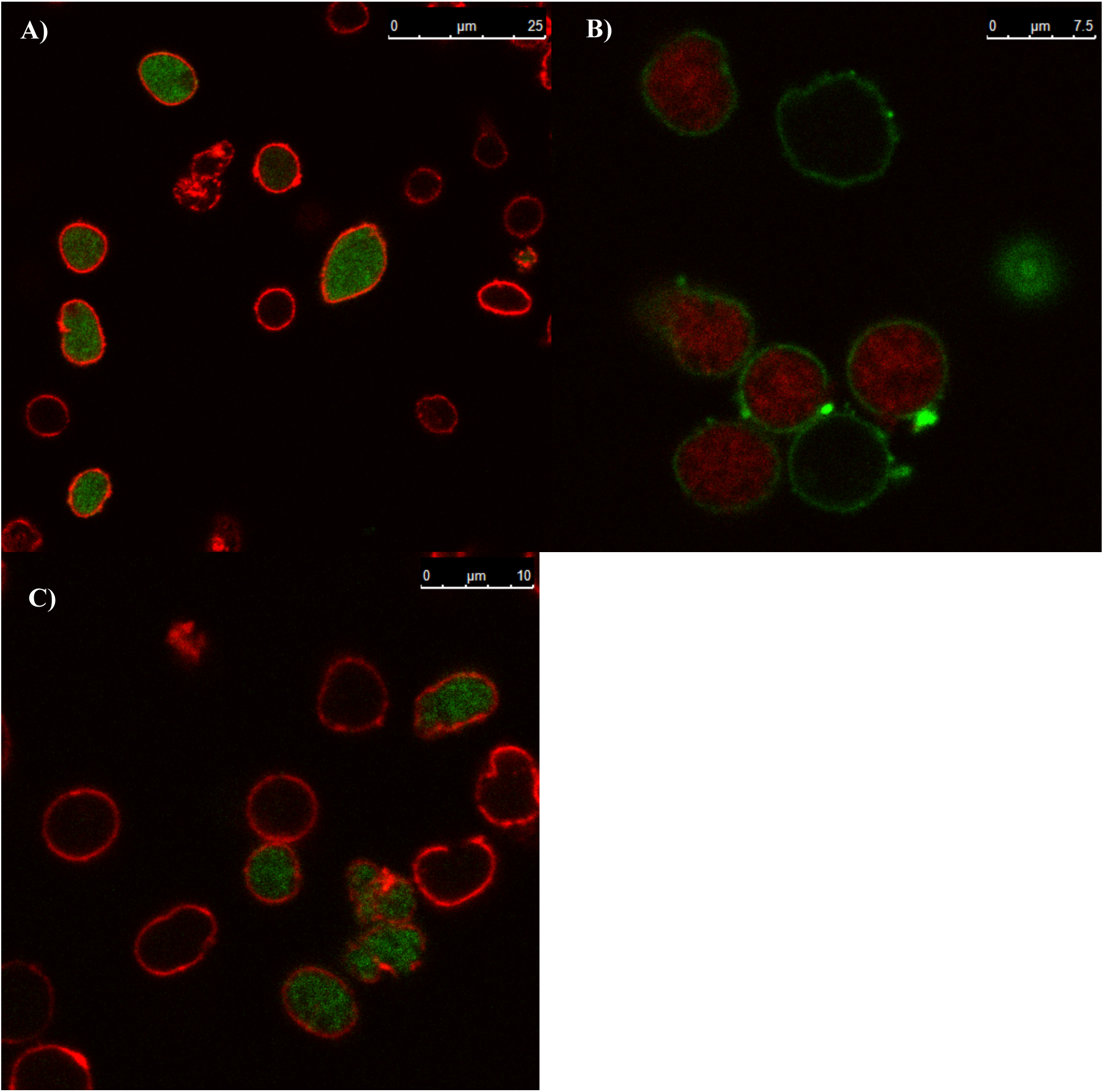
Confocal microscope images of isolated brain nuclei. The nuclear membrane (mCLING) and cell-type specific transcription factors are visualized to evaluate nuclei intactness and FANS sorting markers. Scale of the nuclei is visualized in the upper right corner. **A)** mCLING-647 (red) and NeuN (green), **B)** mCLING (green) and SOX10 (red), **C)** mCLING (red) and IRF5 (green).

**Figure S3.**
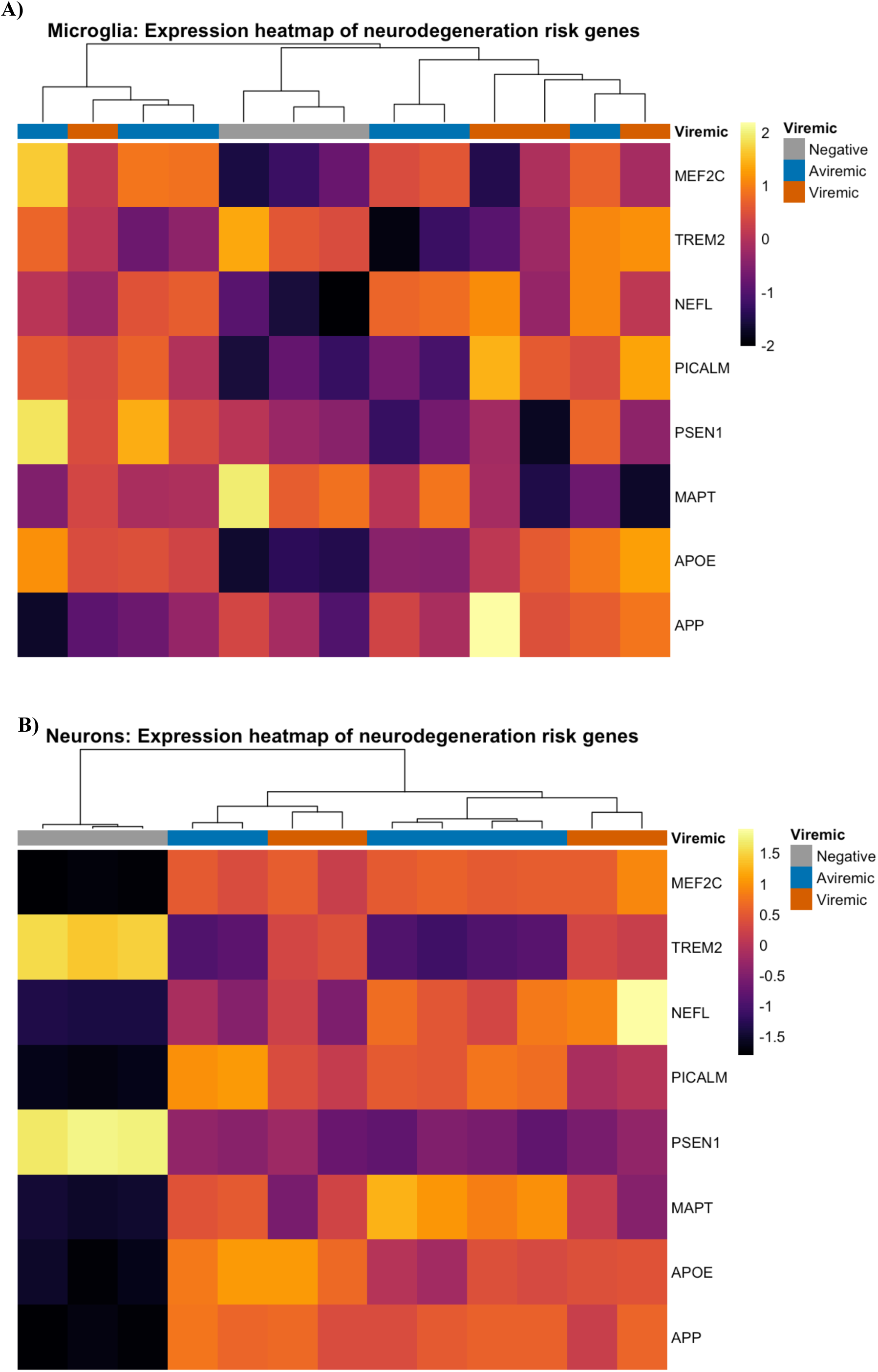
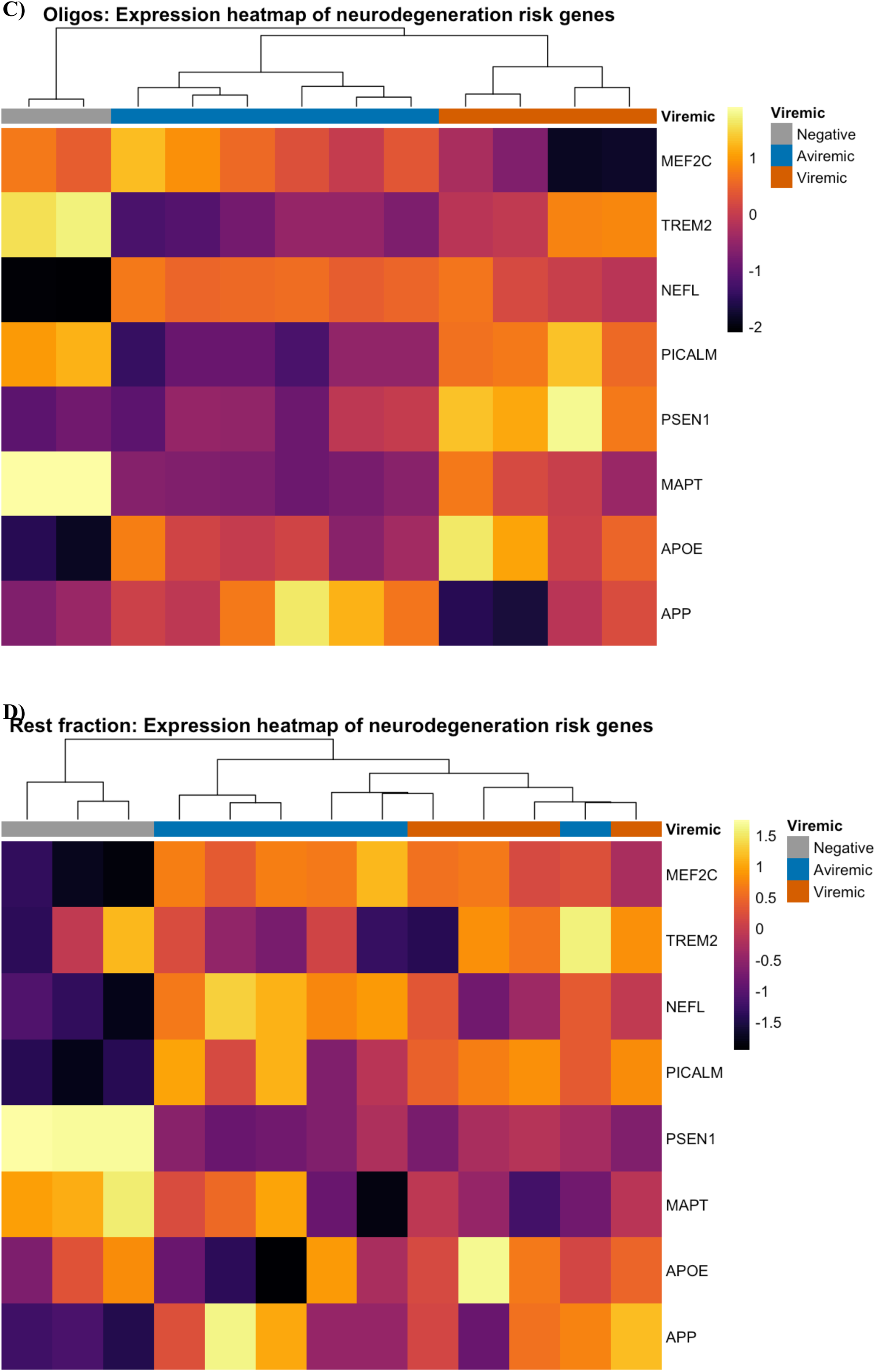
Normalized expression of neurodegeneration markers in the cellular fractions. **A)** Microglia, **B)** Neurons, **C)** Oligodendrocytes, **D)** Rest fraction.

---

**Table S1.**
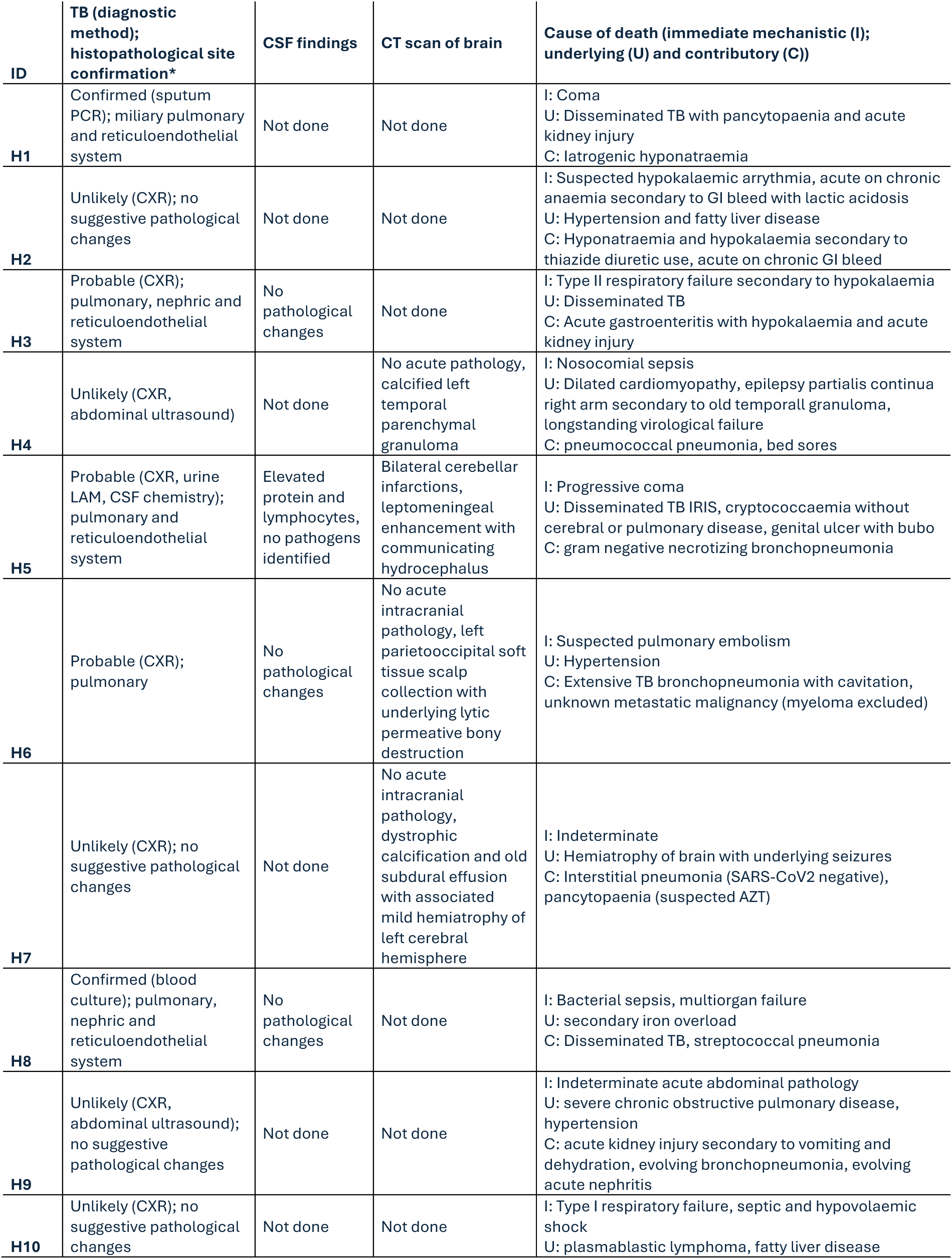
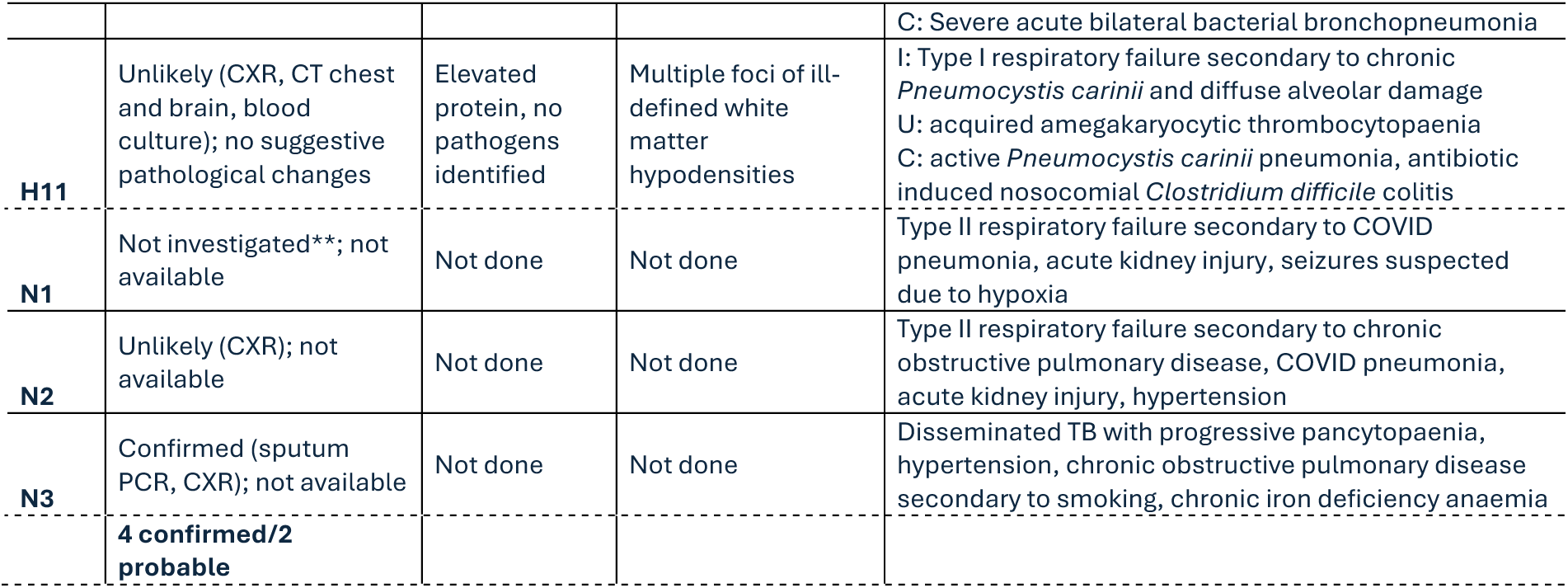
Detailed clinical records and cause of death of the study participants. The ID of individuals with HIV starts with “H”, the HIV-negative individuals start with “N”. CSF = cerebrospinal fluid, CXR= chest-x-ray, LAM=lipoarabinomannan

**Table S2.** Overview of quality of bulk-sequencing data. After filtering on sample level per cell type, the number of genes that is left per subset is indicated in the last column. IQR values are given between brackets.

|  | Median number raw reads (million) | Median mapped reads (percentage) | Median number of unique genes | Number of genes DEG analyses |
| --- | --- | --- | --- | --- |
| **Microglia** | 4.2 [3.1-6.1] | 66.0 [63.3-72.3] | 15,350 [14,375-16,025] | 8,943 |
| **Neurons** | 21.2 [15.2-45.2] | 62.8 [58.1-72.1] | 17,850 [17,325-20,275] | 11,765 |
| **Oligodendrocytes** | 27.5 [13.0-43.3] | 62.9 [55.15-76.9] | 19,200 [17,350-20,075] | 11,832 |
| **Rest fraction** | 13.6 [7.3-19.6] | 68.1 [60.7-76.1] | 18,500 [16,625-19,175] | 10,942 |

**Table S3.**
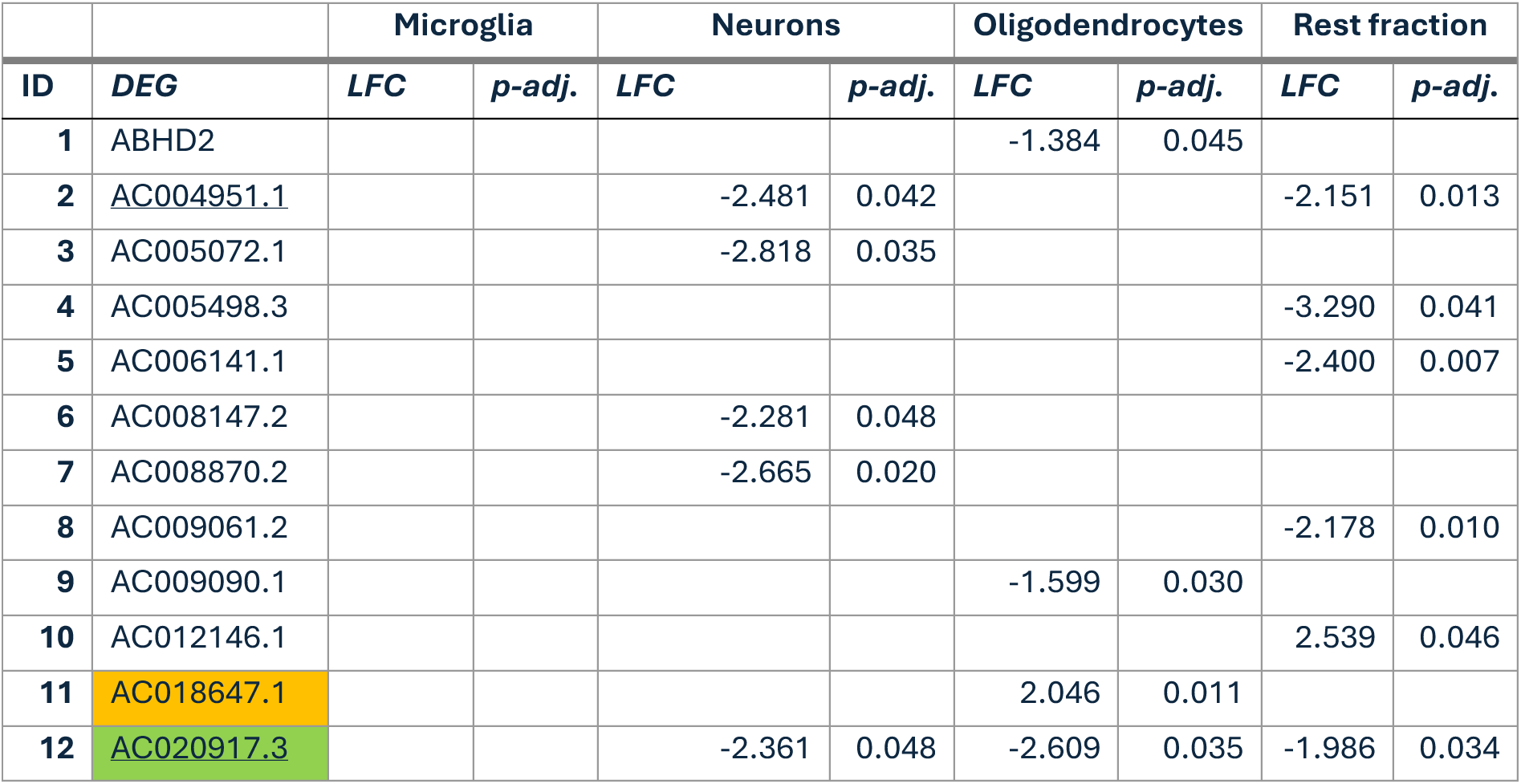
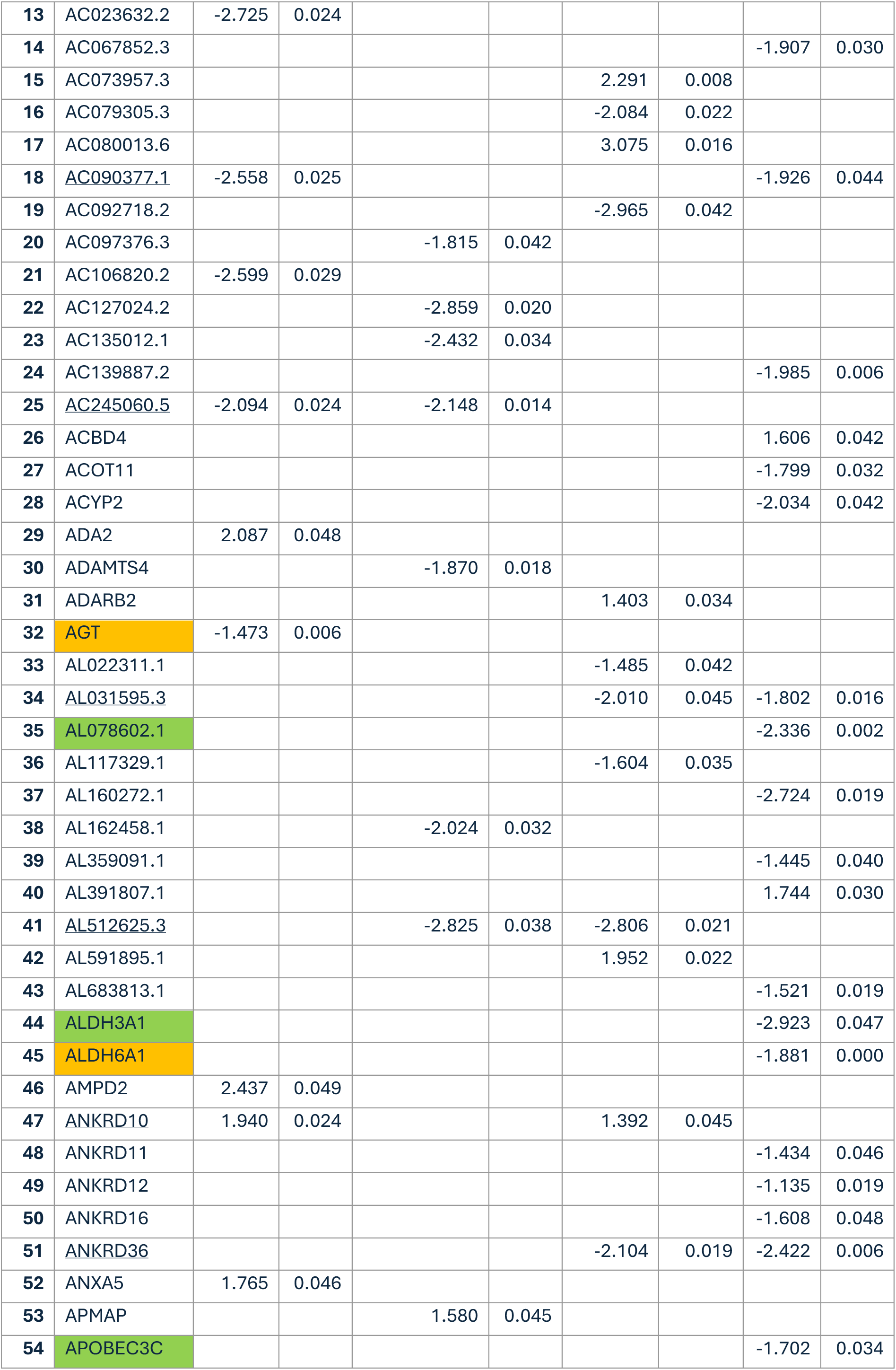
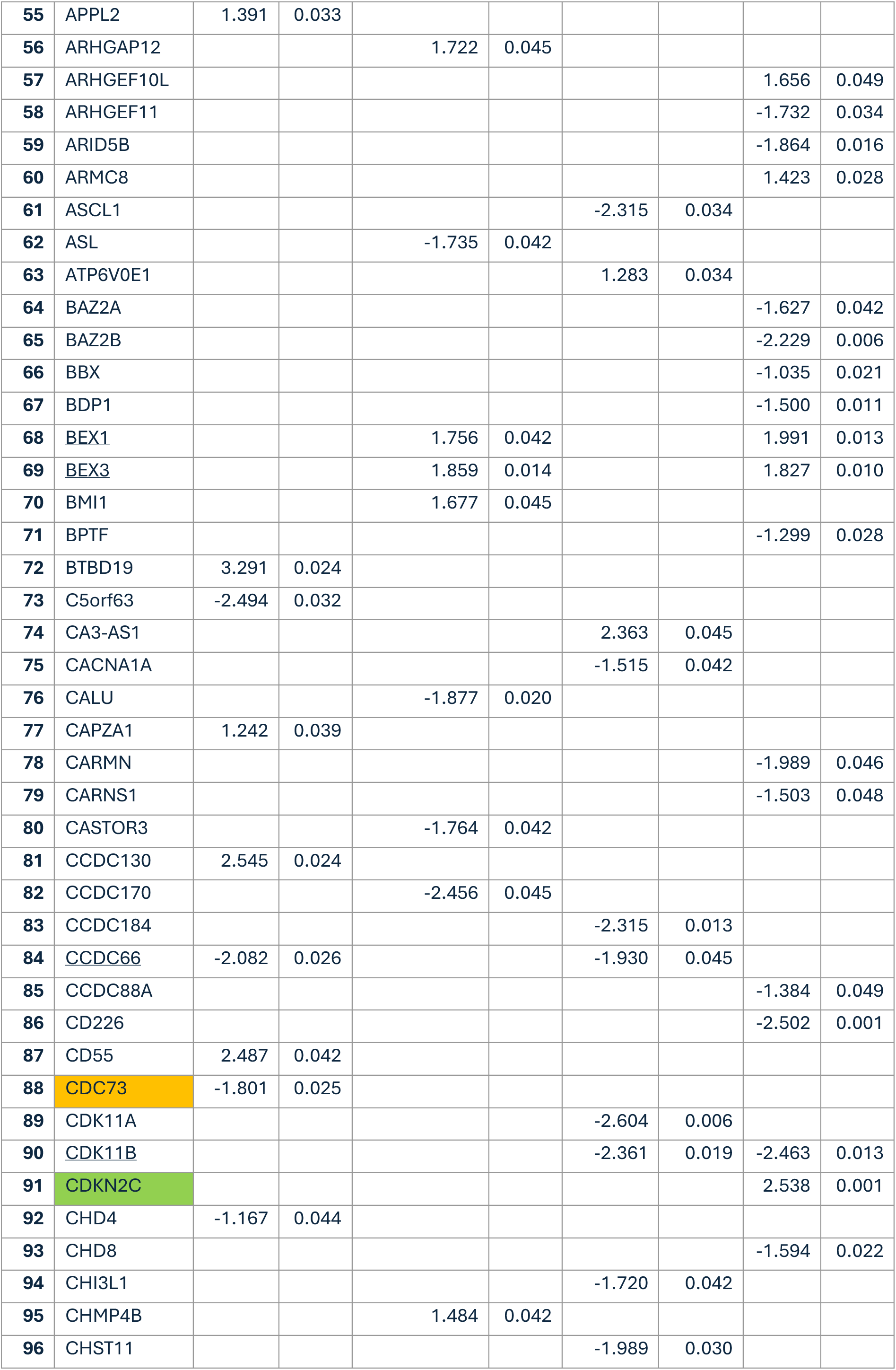
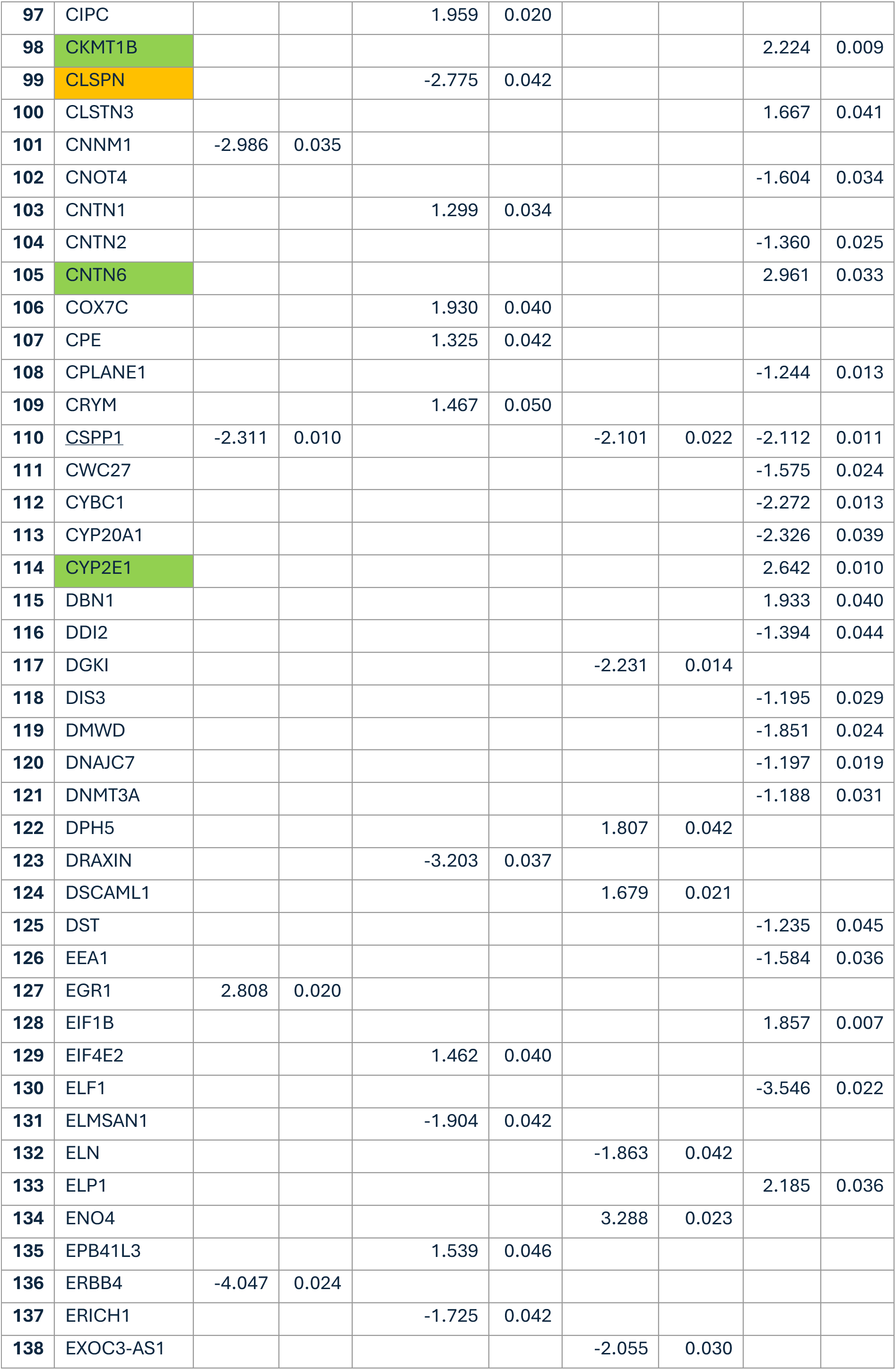
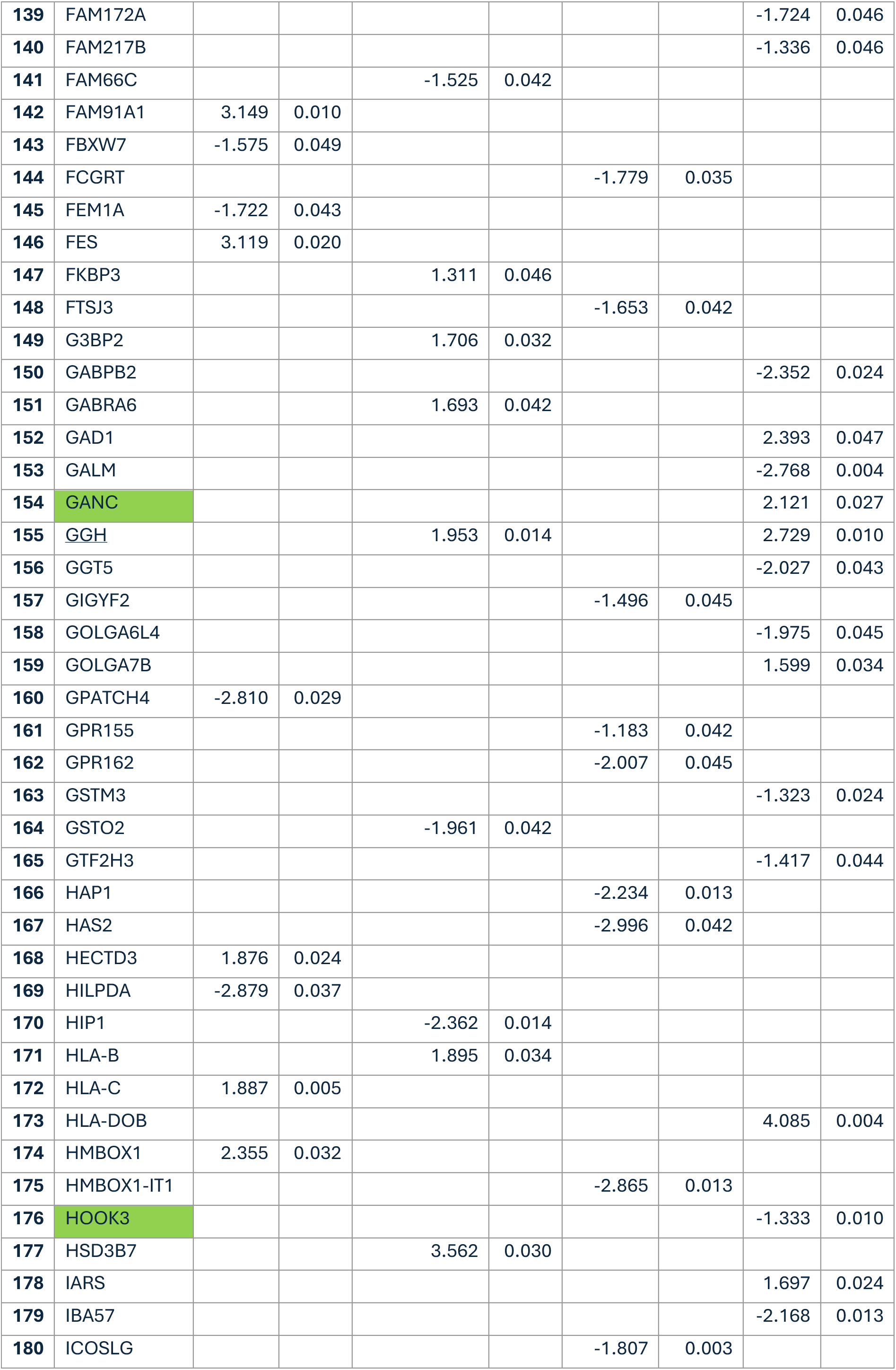
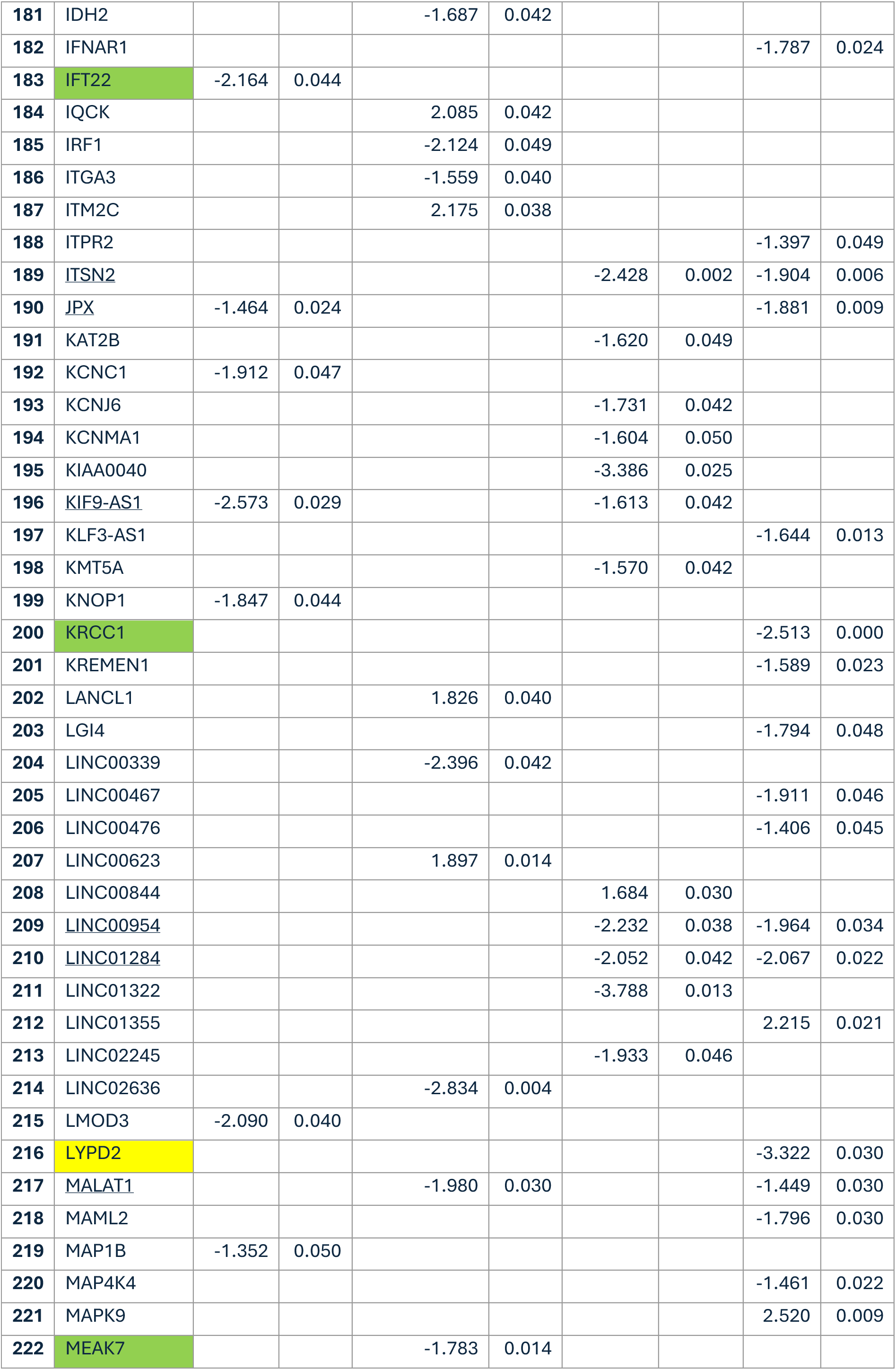
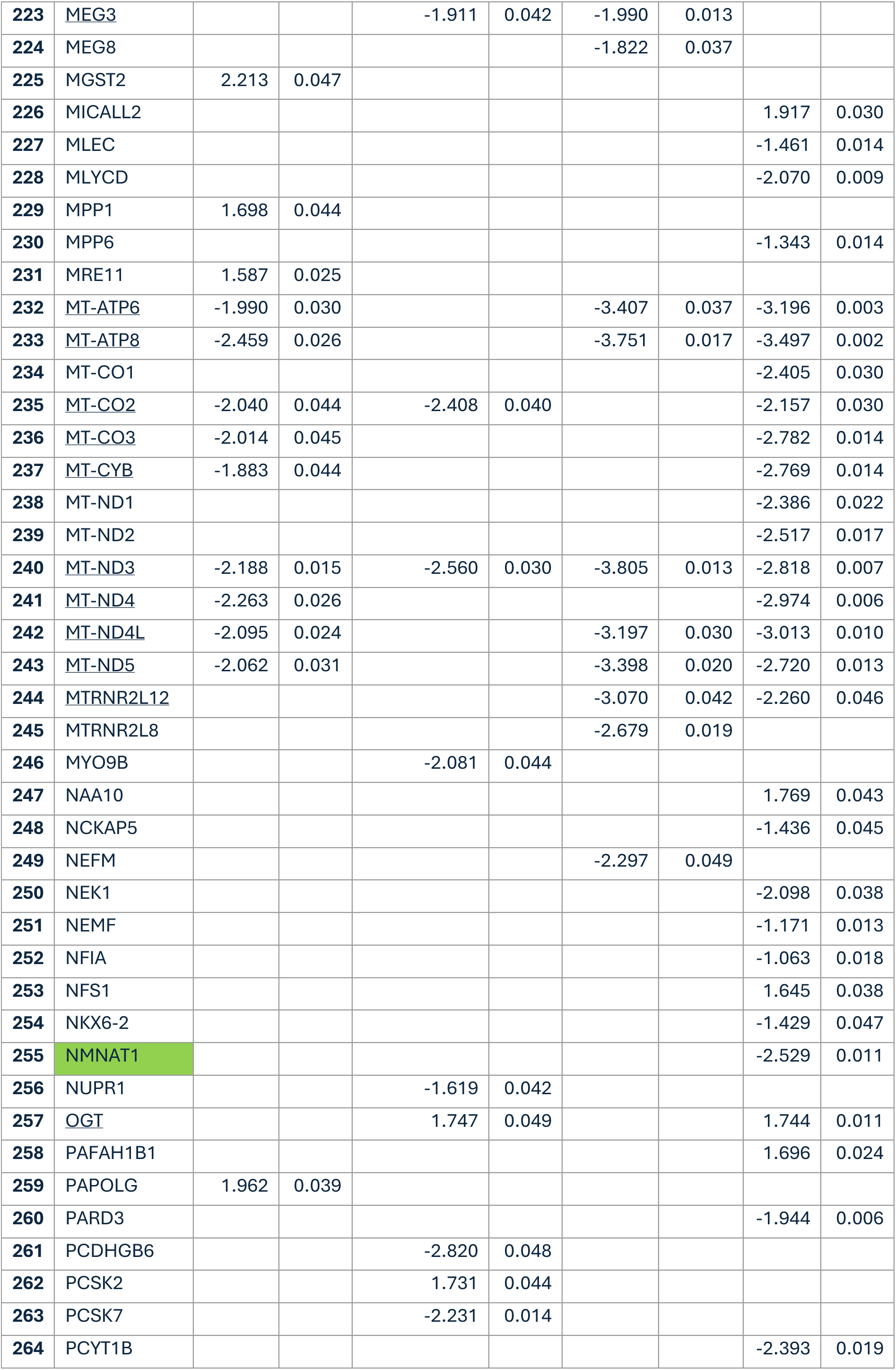
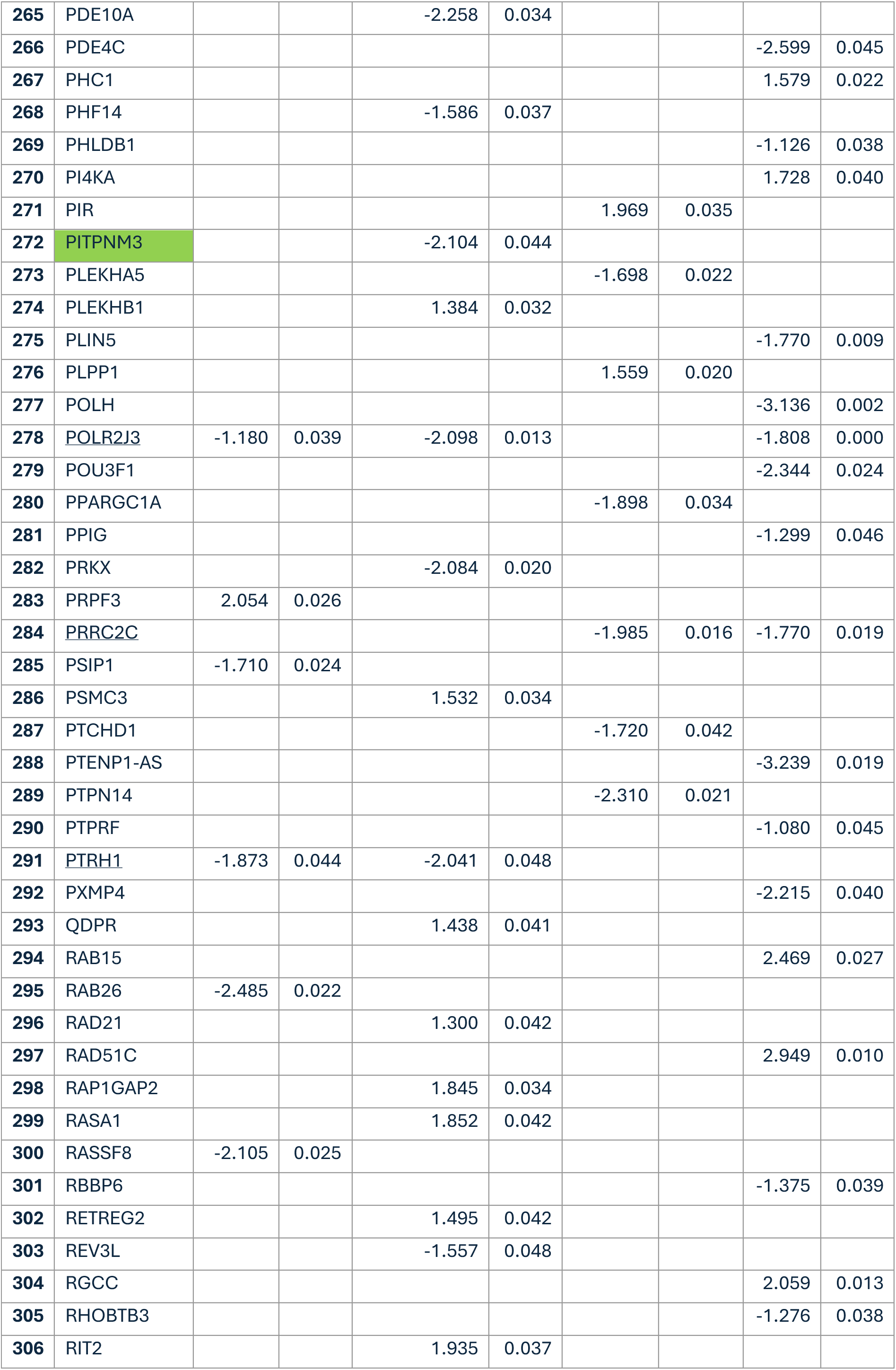
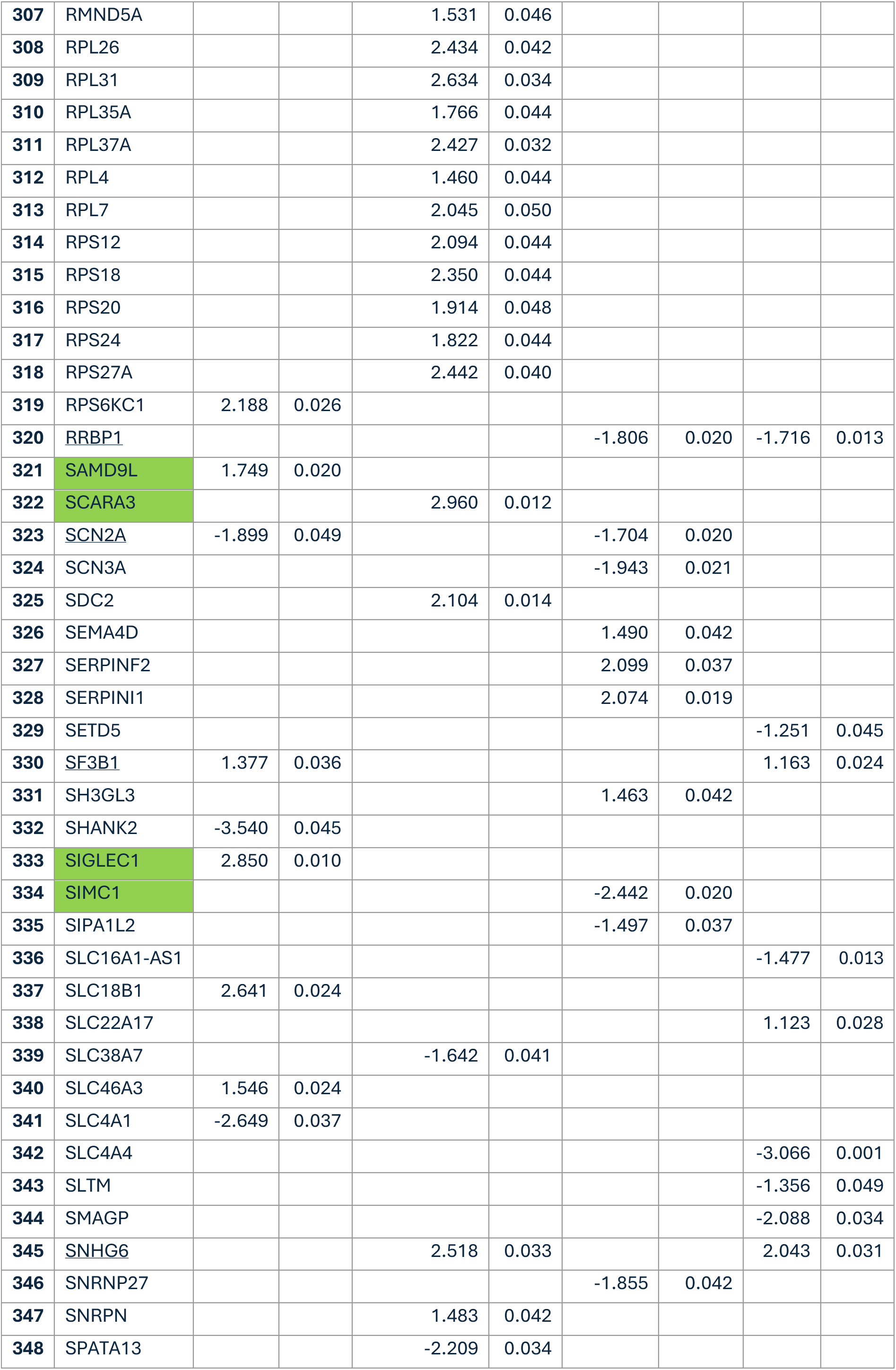
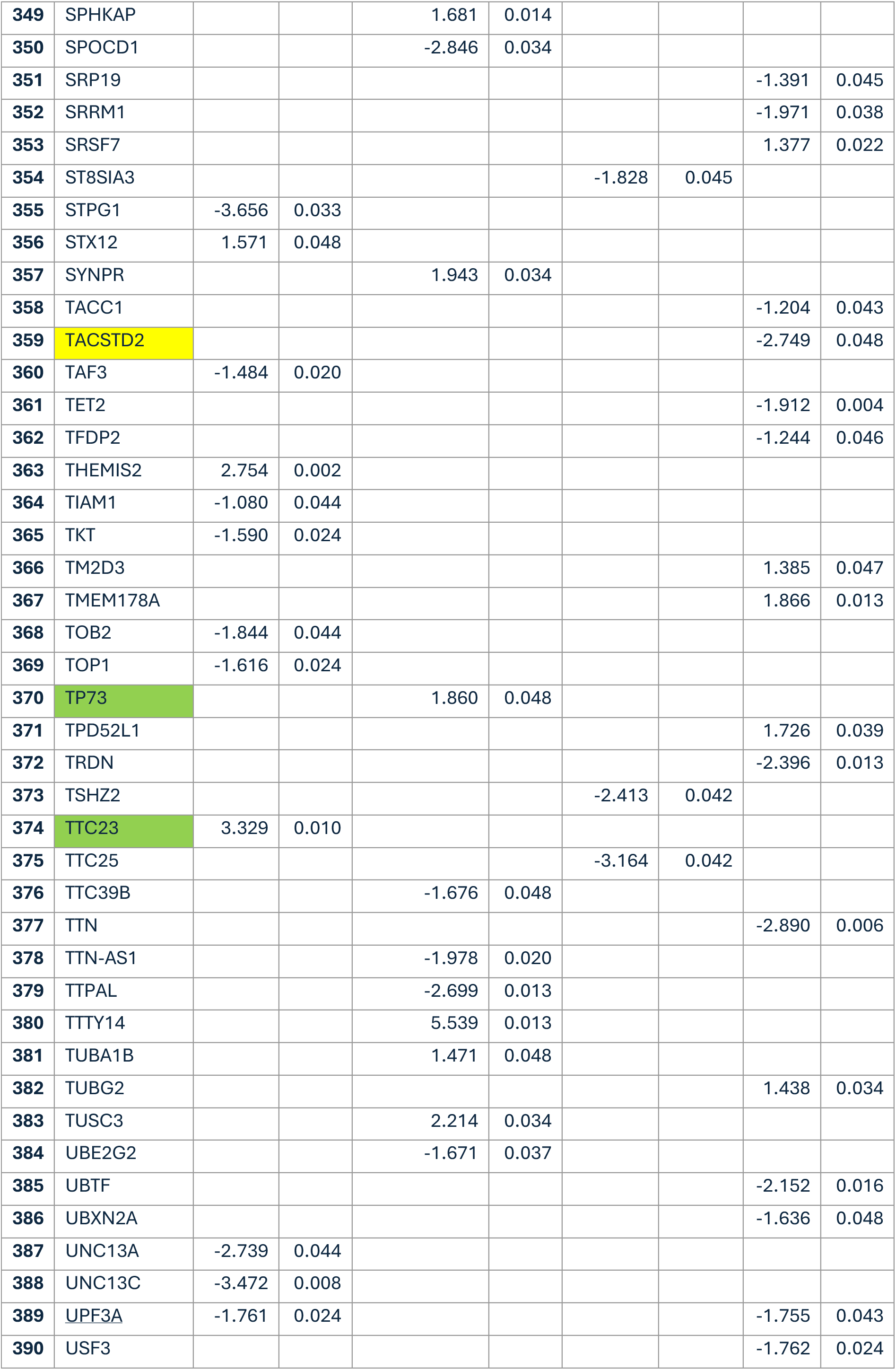
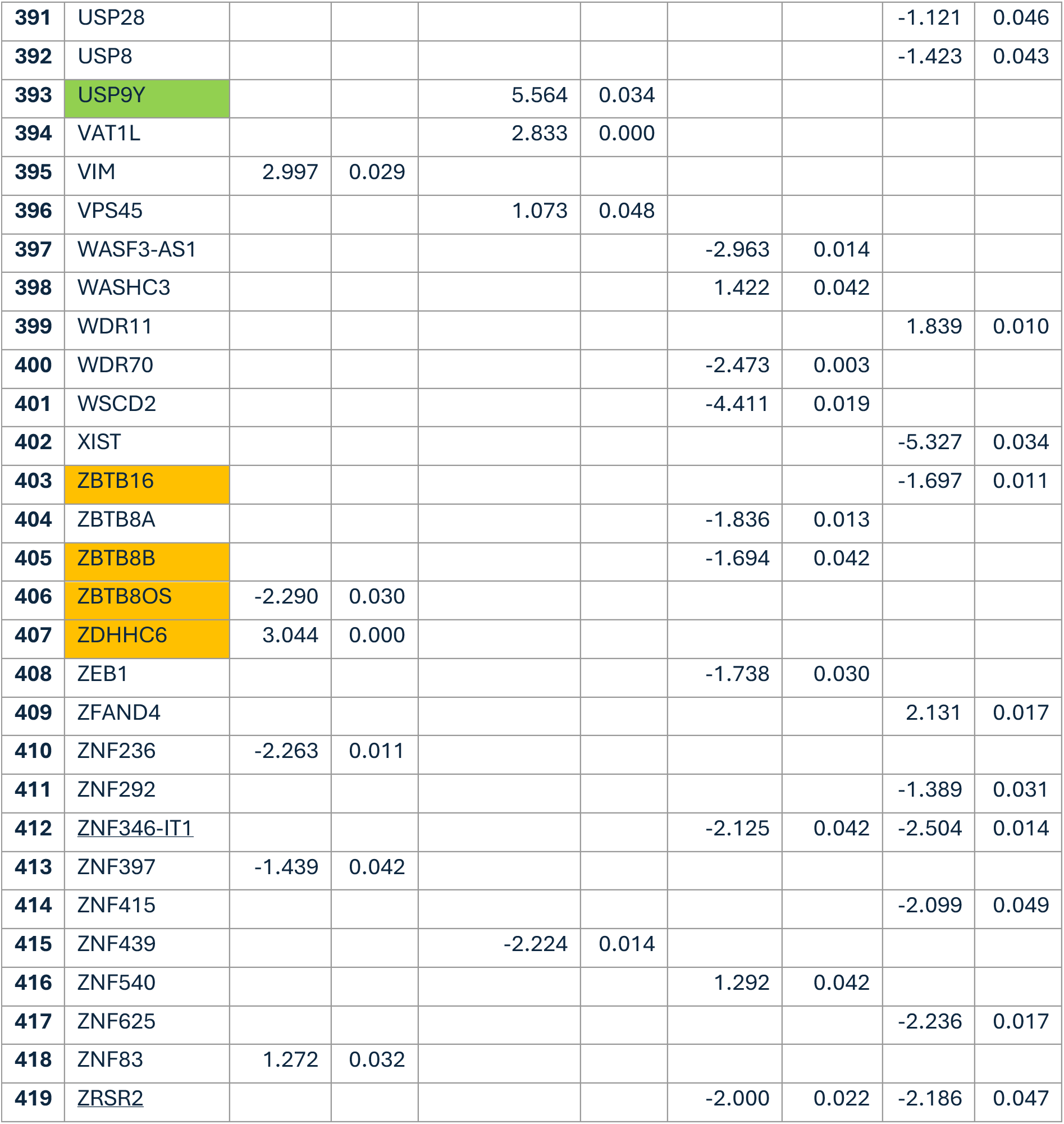
Differently expressed genes per cell type for viremic versus aviremic individuals. Genes differently expressed within more than 1 cellular subset are underlined. Green labeled genes are also DEGs when comparing aviremic individuals and HIV-negative individuals, orange labeled genes are also DEGs when comparing viremic individuals with HIV-negative individuals and yellow labelled genes are DEGs in all 3 comparisons.

**Table S4.** Literature search of DEGs in viremic versus aviremic individuals. Literature research overview of DEGs occurring in multiple cell types with search terms “HIV”, “inflammation” and “infection” or “CNS” in combination with the DEG. A summary of the findings and the associated PMID of the article are given. M = Microglia, N = Neurons, O = Oligodendrocytes, R= Rest fraction

| ID | ID in Table S3 | DEG | Direction | Cell type | Association with | Pubmed ID (PMID) |
| --- | --- | --- | --- | --- | --- | --- |
| 1 | 2 | AC004951.1 | Down | N, R | None | - |
| 2 | 12 | AC020917.3 | Down | N, O, R | None | - |
| 3 | 18 | AC090377.1 | Down | M, R | None | - |
| 4 | 25 | AC245060.5 | Down | M, N | None | - |
| 5 | 34 | AL031595.3 | Down | R, O | None | - |
| 6 | 41 | AL512625.3 | Down | N, O | None | - |
| 7 | **47** | ANKRD10 | Down | M, O | None | - |
| 8 | **51** | ANKRD36 | Down | O, R | Elevated methylation of ANKRD36 in postmortem brain samples of people with HIV | 33805201 |
| 9 | **68** | BEX1 | Up | N, R | Indicated as biomarker for cryptococcosis-associated immune reconstitution inflammatory syndrome in PWH infected with cryptococcal meningitis after ART. | 30038928 |
| 10 | **69** | BEX3 | Up | N, R | Involved in neurological disorders and relevant for higher brain function in placental mammals. | 33100228 |
| 11 | **84** | CCDC66 | Down | M, O | None | - |
| 12 | **90** | CDK11B | Down | O, R | None | - |
| 13 | **110** | CSPP1 | Down | M, O, R | None | - |
| 14 | **155** | GGH | Up | N, R | Indicated in HIV latency reactivation pathways. Upregulated more than 10 folds in bryostatin (LRA) stimulated CD4+ T cells of PWH on successful ART for >36 months and a viral load <50 copies/mL. | 32103135 |
| 15 | **189** | ITSN2 | Down | O, R | None | - |
| 16 | **190** | JPX | Down | M, R | None | - |
| 17 | **196** | KIF9-AS1 | Down | M, O | None | - |
| 18 | **209** | LINC00954 | Down | O, R | None | - |
| 19 | **210** | LINC01284 | Down | O, R | None | - |
| 20 | **217** | MALAT1 | Down | N, R | HSV-2 contributes to HIV-1 reactivation through various mechanisms, including upregulation of MALAT1. MALAT1 was also found to be dramatically upregulated in HIV-1 infected macrophages, where MALAT1 overexpression also enhanced viral replication. | 37079384, 40414289 |
| 21 | **223** | MEG3 | Down | N, O | Inhibits M2 macrophage polarization (anti-inflammatory) in viral myocarditis, dysregulation of MEG3 is associated with several neurodegenerative diseases. | 33047847, 39675694 |
| *Mitochondrial RNA*<br>*(marked in purple, ID 22-31)* |  |  | Down | variable | Brain HIV RNA loads are inversely correlated with genesets representative of oxidative phosphorylation and the TCA cycle in the frontal cortex, basal ganglia and white matter in postmortem tissue of DPWH. Advanced mitochondrial (mt) aging, and accelerated decrease of mtDNA was reported in postmortem brain tissue of DPWH, a change driven by HIV and not associated with ART exposure. Multiple articles report accumulative mtDNA damage and mutations caused by HIV infection and HIV-1 Tat. The collective impairment of mitochondrial complexes I, II, III and IV by the downregulation of MT-ND1, MT-ND2, MT-ND3, MT-ND4L, MT-CYB, MT-CO3 and MT-ATP6 across cell types in postmortem frontal cortex tissue of individuals with COVID-19 was reported by using RNA-seq and snRNA-seq. | 34108514, 32722761, 40494474, 39271627 |
| 22 | **232** | MT-ATP6 | Down | M, O, R | Decreased in people with HIV whose CD4 count does not return to normal levels after start ART. Some mutations in MT-ATP6 and MT-CYB disrupt the protein stability and are relatively common in COVID-19 patients compared to healthy controls. Mutations in MT-ATP6/8 can cause disruption of the ATP synthase complex and result in various clinical features based on brain MRI. See overview. | 36195585, 40839580, 40112238, 39271627 |
| 23 | **233** | MT-ATP8 | Down | M, O, R | See MT-ATP6. | 40112238 |
| 24 | **235** | MT-CO2 | Down | M, N, R | MT-CO2 was previously reported to be unchanged in primary human astrocytes upon treatment with ART *in vitro* in a culture without any HIV+ cells or virus added. | 41369393 |
| 25 | **236** | MT-CO3 | Down | M, R | See overview. | 39271627 |
| 26 | **237** | MT-CYB | Down | M, R | The genomic regions of MT-RNR1, MT-ND5 and MT-CYB were differentially methylated upon treatment with HIV-1 Tat or cocaine in primary human astrocytes. See overview. | 33100130, 39271627 |
| 27 | **240** | MT-ND3 | Down | M, N, R, O | Various mutations are involved in Leigh syndrome (a neurodegenerative disease) and axonal polyneuropathy, decreased in people with HIV whose CD4 count does not return to normal levels after start ART. See overview. | 14764913, 33732874, 36195585 |
| 28 | **241** | MT-ND4 | Down | M, R | See overview. | 39271627 |
| 29 | **242** | MT-ND4L | Down | M, O, R | See overview. | 39271627 |
| 30 | **243** | MT-ND5 | Down | M, O, R | MT-ND1, MT-ND5 and MTRNR2L12 are protein coding genes related to NADH dehydrogenase activity and the apoptotic process, and were differentially expressed and downregulated in platelets of COVID-19 patients compared to healthy controls. | 35421970 |
| 31 | **244** | MTRNR2L12 | Down | O, R | See MT-ND5. | 35421970 |
| 32 | **257** | OGT | Up | N, R | Overexpression of OGT and the subsequent O-GlcAcylation of transcription factor Sp1 are reported to inhibit the HIV-1 LTR in primary CD4+ lymphocytes. OGT plays an important role in immune-metabolic crosstalk, also in the CNS, and responds upon cellular stressors to reduce immune inflammation. | 19193796, 30770249, 30770249, 36757814 |
| 33 | **278** | POLR2J3 | Down | M, N, R | Association with infected resting CD4 T cells | 38156317 |
| 34 | **284** | PRRC2C | Down | O, R | None | - |
| 35 | **291** | PTRH1 | Down | M, N | None | - |
| 36 | **320** | RRBP1 | Down | O, R | Downregulated in CD4 latency model and known for its interaction with Vpr | 36090997 |
| 37 | **323** | SCN2A | Down | M, O | Implied in seizure disorders, epilepsy and (epileptic) encephalopathy. Dysregulation of the FOXD3/SCN2A pathway was reported to be promoted through inhibition by PVT1, resulting in neuronal cell injury and neuroinflammation. | 27830185, 12610651, 34156984, 36378391 |
| 38 | **330** | SF3B1 | Up | M, O | Interacts with HIV-1 Tat. Inhibition of SF3B1 prevented HIV reactivation from latency irrespective of the used LRA in primary CD4+ T cells, monocyte-derived macrophages and Jurkat, THP-1 and U87 cell lines. | 30401776 |
| 39 | **345** | SNHG6 | Up | N, R | Expression was significantly lower in PBMCs of COVID-19 patients. | 34147082 |
| 40 | **389** | UPF3A | Down | M, R | UPF3A is a negative regulator of genomic HIV-1 viral RNA nuclear export, and was reported to be excluded from the UFP1-Rev-CRM1-DDX3 nuclear export ribonucleoprotein complex. | 26492277 |
| 41 | **412** | ZNF346-IT1 | Down | O, R | None | - |
| 42 | **419** | ZRSR2 | Down | O, R | None | - |

**Table S5.**
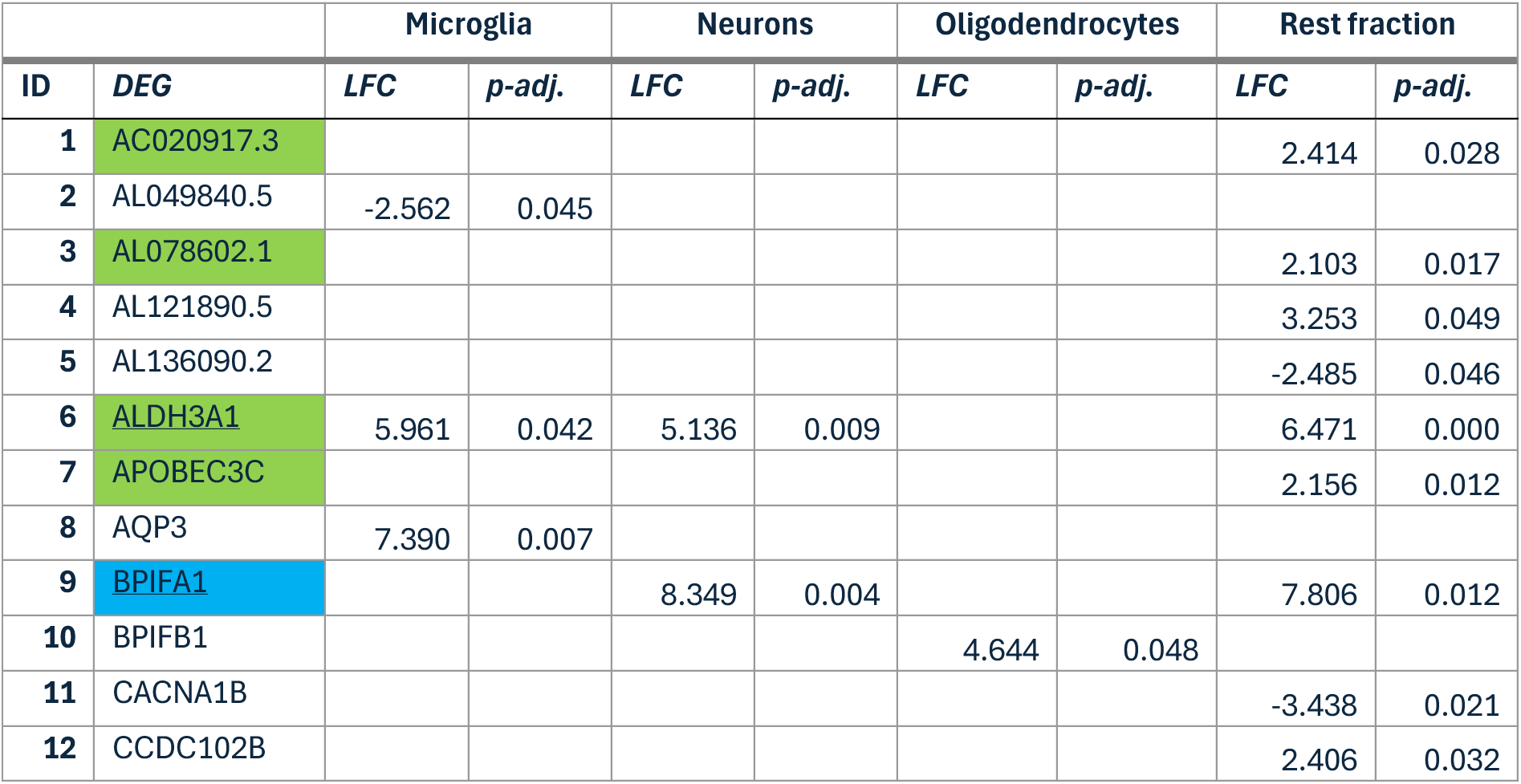
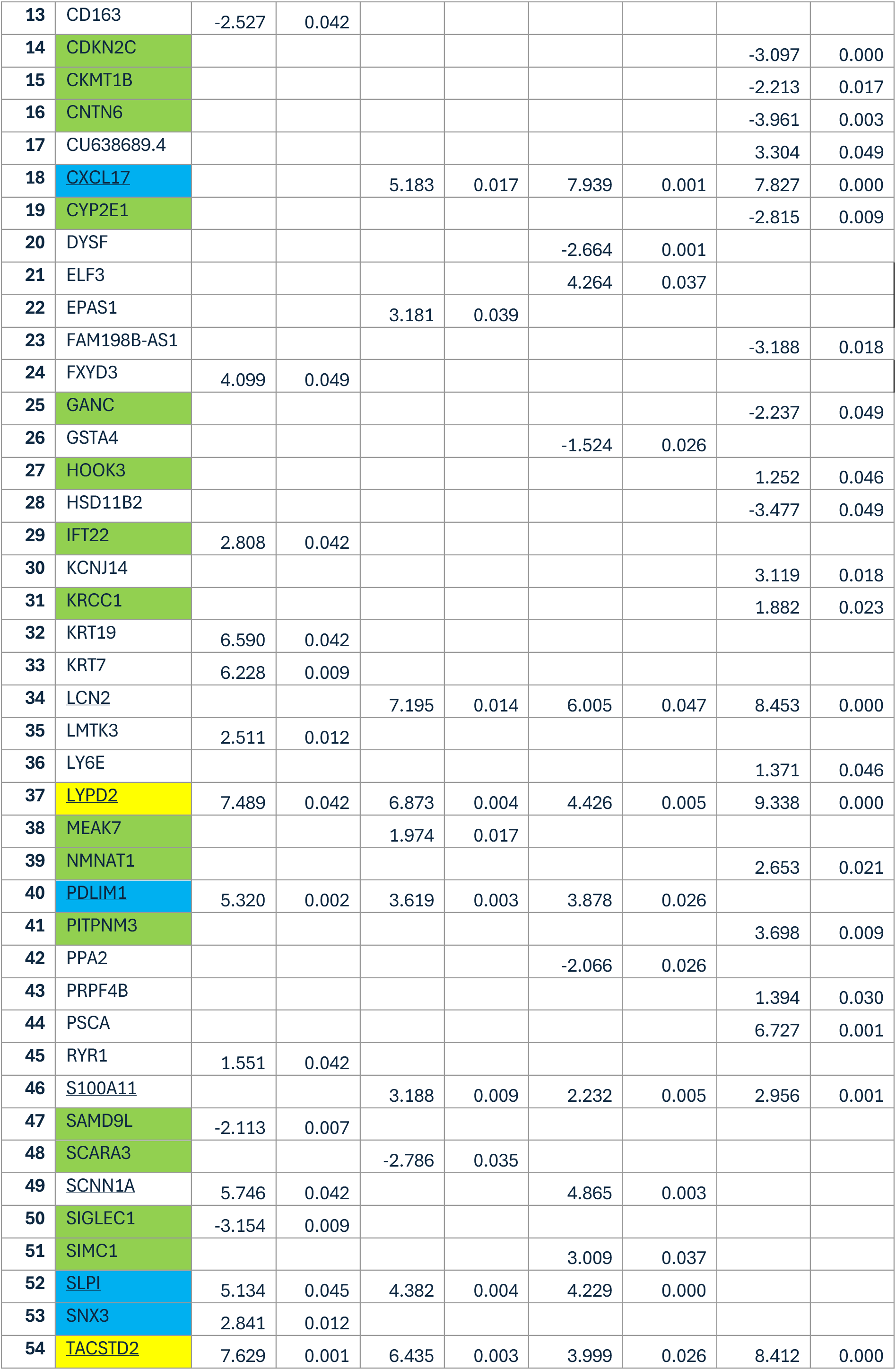
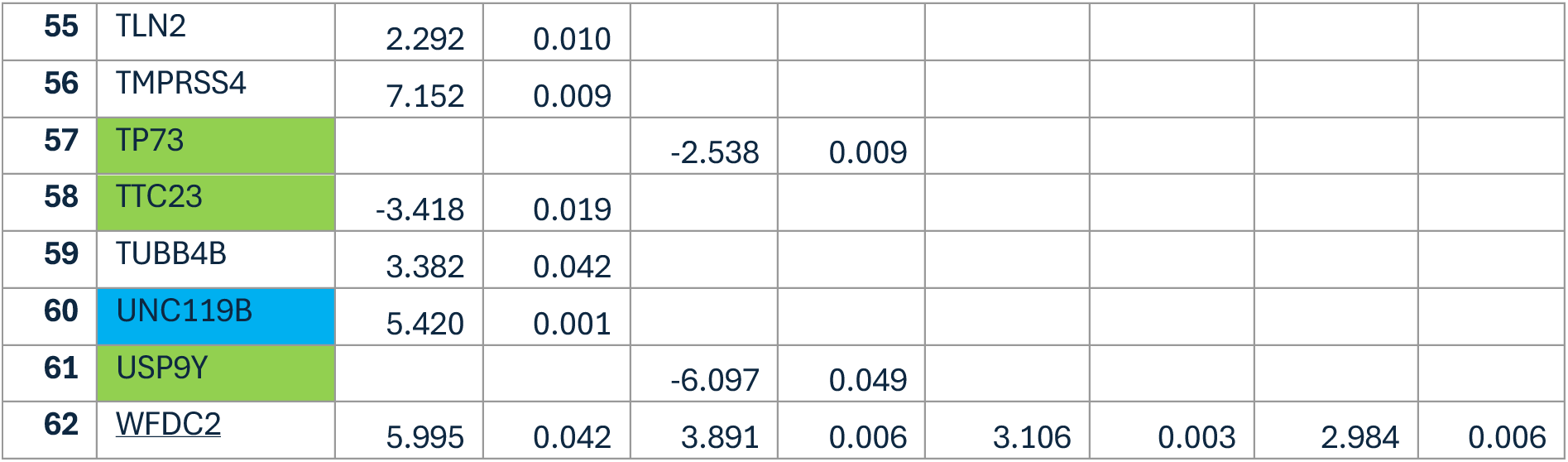
Differently expressed genes per cell type for aviremic individuals versus HIV-negative individuals. Genes differently expressed within more than 1 cellular subset are underlined Blue labeled genes are also DEGs when comparing viremic and aviremic, green labeled genes are also DEGs when comparing viremic individuals with HIV-negative individuals and yellow labelled genes are DEGs in all 3 comparisons.

**Table S6.**
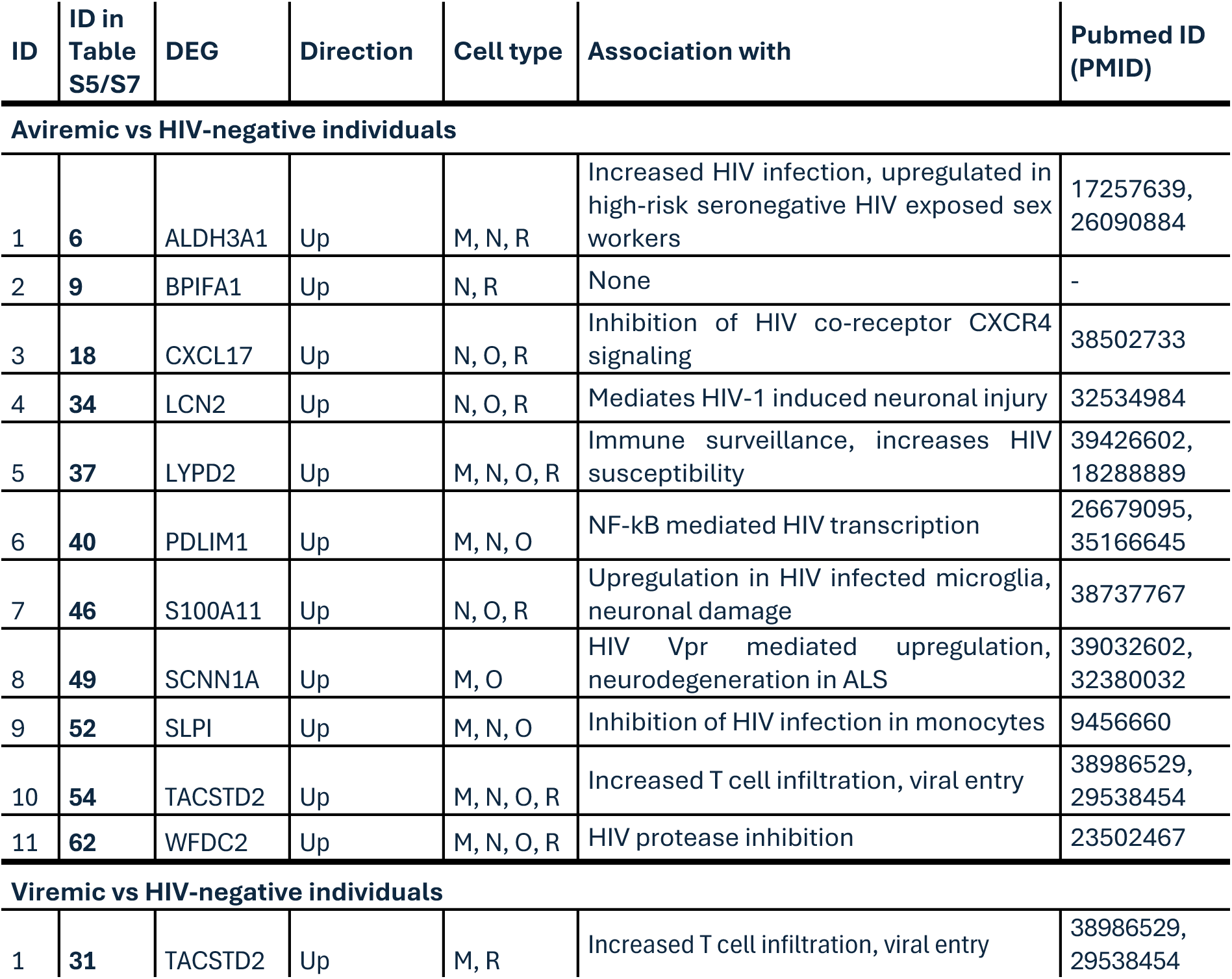
Literature search of DEGs in viremic or aviremic individuals versus HIV-negative individuals. Literature research overview of DEGs occurring in multiple cell types with search terms “HIV”, “inflammation” and “infection” or “CNS” in combination with the DEG. A summary of the findings and the associated PMID of the article are given. M = Microglia, N = Neurons, O = Oligodendrocytes, R= Rest fraction

**Table S7.**
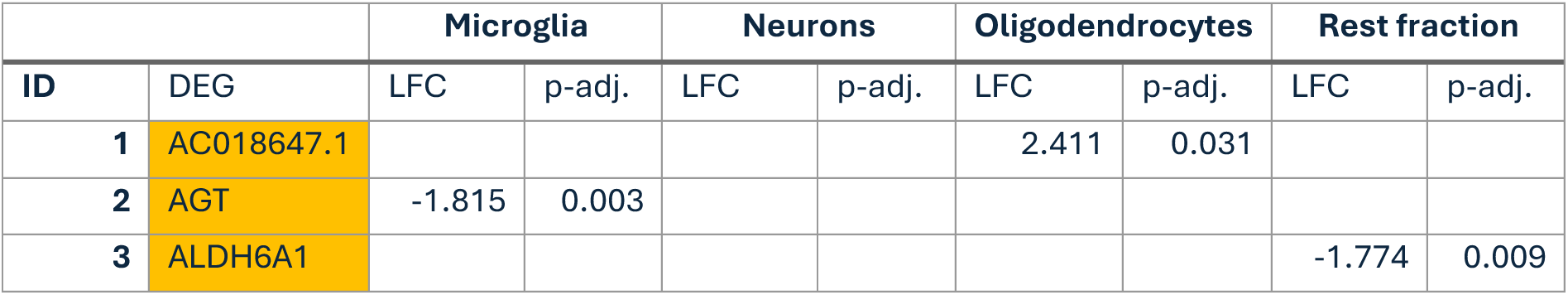
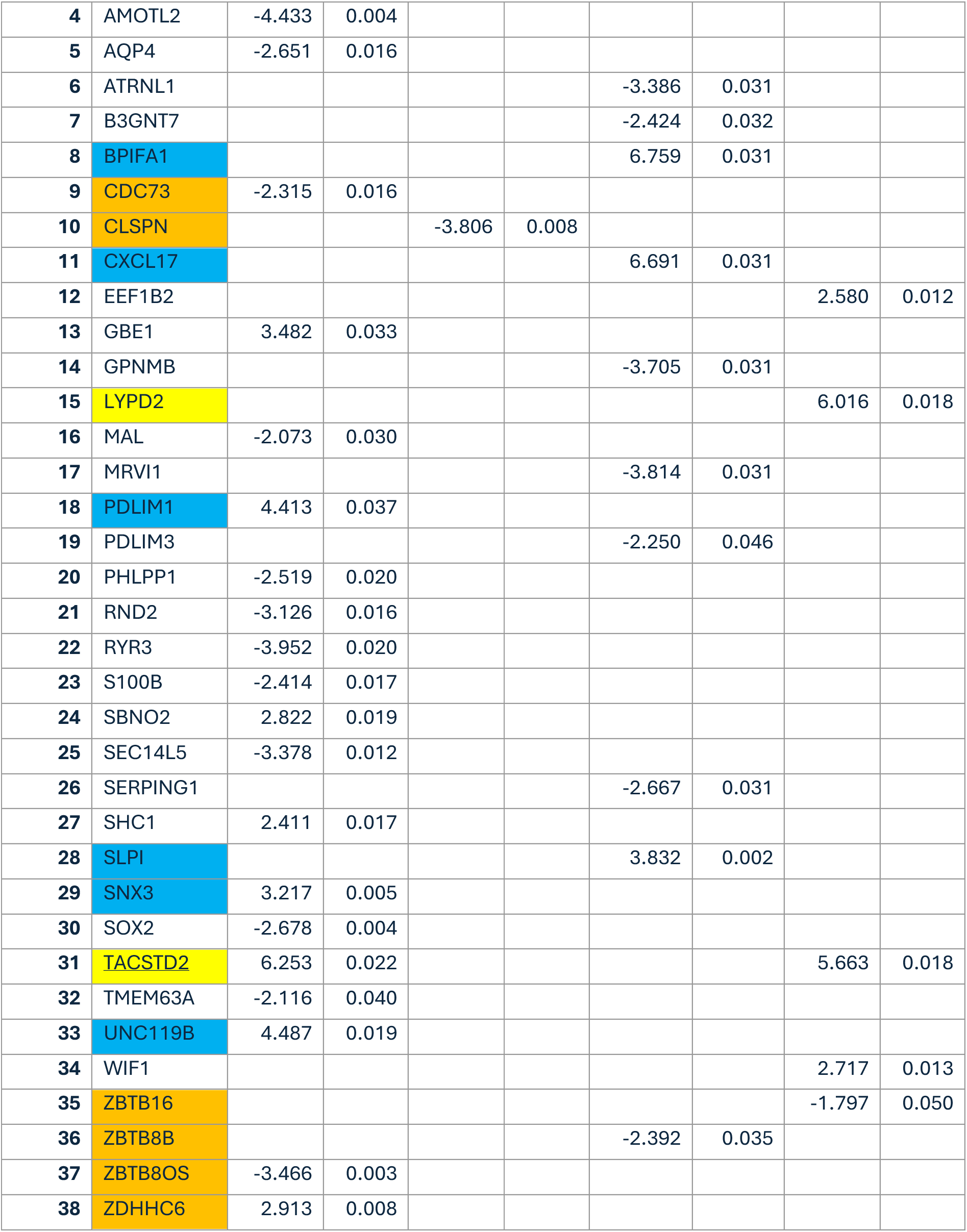
Differently expressed genes per cell type for viremic individuals versus HIV-negative individuals. Genes differently expressed within more than 1 cellular subset are underlined. Blue labeled genes are also DEGs when comparing aviremic individuals with the HIV-negative individuals, orange labeled genes are also DEGs when comparing viremic and aviremic individuals and yellow labelled genes are DEGs in all 3 comparisons.

